# Identification of cell-type-specific response to silicon treatment in soybean leaves through single nucleus RNA-sequencing

**DOI:** 10.1101/2024.04.01.587592

**Authors:** Vikas Devkar, Leonidas D’Agostino, Arjun Ojha Kshetry, Lenin Yong, Altafhusain B Nadaf, VP Thirumalaikumar, Aleksandra Skirycz, Jianxin Ma, Robert M. Stupar, Luis Herrera-Estrella, Rupesh Deshmukh, Gunvant B. Patil

**Affiliations:** Institute of Genomics for Crop Abiotic Stress Tolerance, Department Plant and Soil Sciences, Texas Tech University, Lubbock, TX 79409, USA; Department of Botany, Savitribai Phule Pune University, Pune, MH 411007, India; Proteomics facility, Bindley Bioscience Center, Purdue University, West Lafayette, IN 47907, USA; Department of Biochemistry & Molecular Biology, Michigan State University, East Lansing, MI 48824, USA; Department of Agronomy and Center for Plant Biology, Purdue University, West Lafayette, IN 47907, USA; Department of Agronomy and Plant Genetics, University of Minnesota, St. Paul, MN 55108, USA; Department of Biotechnology, Central University of Haryana, Mahendragarh, HR 123031, India

## Abstract

In agriculture, mineral nutrients uptake and deposition profoundly influence plant development, stress resilience, and productivity. Despite its classification as a non-essential element, silicon (Si) is crucial in plant physiology, particularly in defense response and stress mitigation. While genetic and molecular mechanisms of Si uptake and transport are well-studied in monocots, particularly rice, its role in dicot species, such as soybean, remains unclear at the cellular and molecular levels. Traditional bulk transcriptomics methods lack the resolution to uncover cellular heterogeneity. Here, we present a study by utilizing single-nucleus RNA sequencing (snRNA-seq) to dissect cellular responses to Si accumulation in soybean leaves. Our analysis revealed distinct cellular populations, including a novel Si-induced cell cluster within vascular cells, suggesting a specific mechanism of Si distribution. Si treatment induced the expression of defense-related genes, particularly enriched in vascular cells, highlighting their specialized role in activating plant defense mechanisms. Moreover, Si modulated the expression of genes involved in RNA silencing, phytoalexin biosynthesis, and immune receptor signaling, suggesting a mechanism of transcriptional priming of genes involved in defense responses. We further investigated putative Si transporters, revealing differential expression patterns in response to Si treatment, suggesting presence of active and gradient-based transport mechanisms. Our findings shed light on the vital biotic stress regulatory networks governed by Si treatment in soybean leaves, paving potential strategies for enhancing stress tolerance and agronomic performance in crops.

## Introduction

Mineral nutrients are essential elements that plants obtain from the soil and use for various physiological processes and are vital for the growth, development, and overall functioning of plants. Silicon (Si) is a lesser-known mineral nutrient but benefits plant health, environmental resilience, and agricultural productivity (Mitani-Uneo et al., 2023). Silicon is not classified as an essential nutrient for plants, Si has been recognized as quasi-essential for its beneficial effects on plant growth and stress tolerance (Coskun et al., 2019; Thakral et al., 2024). Although Si ranks as the earth’s second most abundant element in the earth crust, the role of Si in plant development, abiotic stress tolerance, and disease resistance was overlooked until the discovery of Si transporter genes in rice (Ma et al., 2006). Silicon gets absorbed through the plant root system as silicic acid (Si (OH)_4_) and transported to aerial tissues. The deposition of Si in growing tissue forms a complex with organic compounds and thought to create an apoplastic barrier for invading pathogens (Coskun et al., 2019). These stronger epidermal layers with Si shield plants from environmental stresses and increase resistance to pathogens. Recent evidences based on comparative genomics, physiological, and molecular studies showed that Si uptake improves resistance against several fungal diseases such as powdery mildew (Wang et al., 2020), rice-blast (Sathe et al., 2021), stem rust (Li et al., 2022), root-rot (Abbai et al., 2019), frog-eye disease (Telles et al., 2014), and Phytophthora (Rasoolizadeh et al., 2018) in various economically important crops. Silicon also plays a beneficial role in abiotic stresses, including salt and osmotic stress tolerance in crops like cucumber and sorghum and reducing heavy metal toxicity in rice, wheat, and maize (reviewed by Thakral et al., 2021).

Polymerization of Si in epidermal layers acts as a modulator influencing the timing and extent of plant defense response. For example, it obstructs pathogenesis-related events, interferes with pathogen signaling, and prevents penetration by pathogenic fungi hypha into the cells (Coskun et al., 2019). Similarly, this Si generated apoplastic barrier restricts evapotranspiration, positively impacting water stress (Thorne et al., 2020). Remarkably, Si uptake, transport, and deposition in various plant species are dynamic and differential processes. In planta, Si content varies from 0.1% to 10% on a dry weight basis (Hodson et al., 2005). Furthermore, in monocots Si deposition occurs in specialized dumbbell shaped cells named silica cells. This mechanism has been extensively studied in rice and sorghum (Ma et al., 2006; Kumar et al., 2017), while in the case of dicots (e.g., soybean and cucumber), Si is deposits in patches on leaf surfaces in between cuticle and epidermal cells (Samuels et al., 1991; Arsenault-Labrecque et al., 2012; Dhingra et al., 2024). Silicon transport mechanisms were very well studied in monocot species. Silicon uptake in rice is mediated by two types of Si transporters, Lsi1 (influx) and, Lsi2 and Lsi3 (efflux) transporters, and shows different polarity in the exodermis and endodermis of root tissue (Ma et al., 2006; Ma et al., 2007; Mitani-Ueno et al., 2023). However, the molecular mechanisms involved in Si uptake, transport, deposition, and its effect on molecular cascades in dicot species, including soybean, have not been very well studied. Moreover, previous studies have primarily relied on bulk transcriptomics, which fails to report for the cellular heterogeneity and does not consider the specific cell types possessing the ability of nutrient uptake and deposition (Hao et al., 2021). The single nucleus RNA-sequencing (snRNA-seq) has enabled researchers to identify cell-specific gene expression in several organisms (Kim et al., 2021; Cervantes-Pérez et al., 2022; Shahan et al., 2022; Tenorio Berrío et al., 2022; Xia et al., 2022; Nobori et al., 2023; Sun et al., 2023).

In the present study, we employed SnRNA-seq approach to identify the cellular heterogeneity of soybean leaf upon Si treatment. We identified GmHiSil2c as an active efflux Si transporter in soybean. We discuss the transformative impact of SnRNA-seq on understanding cellular and molecular networks and biological processes, highlighting the role of Si on plant immunity and uncovering comprehensive transcriptomes of 12 distinct cell types in Si treatment. The analysis identifies specific gene expression patterns related to defense response, immune receptors, RNAi silencing machinery, salicylic acid metabolism, and other processes induced by Si treatment. Additionally, our study discovers a new subset of cells present in higher fractions in Si-treated plants, revealing the dynamic response of plants to Si treatment and providing insights into the subcellular localization of specific Si transporters.

## Result

### SnRNA-seq analysis of Si+ and Si-treated soybean leaves

Soybean plants grown hydroponically and treated with Si (Si+) and without Si (Si-, control) were used to determine Si content in leaf tissues. Following Si+ treatment, Si deposition in the leaves was quantified using scanning electron microscopy, energy-dispersive X-ray spectroscopy (SEM-EDS), and a portable X-ray fluorescence analyzer (pXRF). As expected, Si deposition was observed in Si+ leaves with an average concentration of 5,200 PPM (0.52%) compared to undetectable accumulation in Si-tissues (Figure 1A, and Supplementary Fig. S1). Notably, Si deposition in soybean leaf tissues was not limited to specific cells but rather deposited in patches between epidermal cells and cuticle layer and also in trichomes (Figure 1A and Supplementary Fig. S2). Following high-quality nuclei isolation from control (Si-) and Si-treated (Si+) soybean leaves, nuclei suspension was processed to generate SnRNA-seq libraries using Chromium technology (10X^TM^ Genomics). Illumina sequencing platforms were used to sequence the SnRNA-seq libraries. After pre-filtering, we obtained an average of 2,080 high-quality nuclei with a mean number of 1,547 unique molecular identifiers (UMIs) and 1,007 genes expressed per nucleus (Figure S3A). After employing the Uniform Manifold Approximation and Projection (UMAP) for dimension reduction, 12 distinct cell types were identified based on their marker transcriptomic profiles, broadly classified into three major cell types, namely epidermal, vascular, and mesophyll (Figure 1B, and Supplementary Fig. S3). The percentage analysis of different cell types showed disproportionate distribution of cell types between control and Si+ treatment, especially for clusters #5 and #10 (vascular cell types) and clusters #3 and #7 (mesophyll cells) (Figure 1C). Upset plot analysis revealed each cell type’s contribution to genetic diversity, with palisade mesophyll cells exhibiting the highest portion of uniquely expressed genes, followed by epidermis, spongy mesophyll, and phloem companion cells. Additionally, upset plot analysis exhibited a greater number of unique genes expressed in mesophyll cell cluster (palisade and spongy) followed by vascular (xylem, phloem, companion, vascular bundle), epidermal, Si-induced and guard cells cluster (Figure 1D).

**Figure 1.**
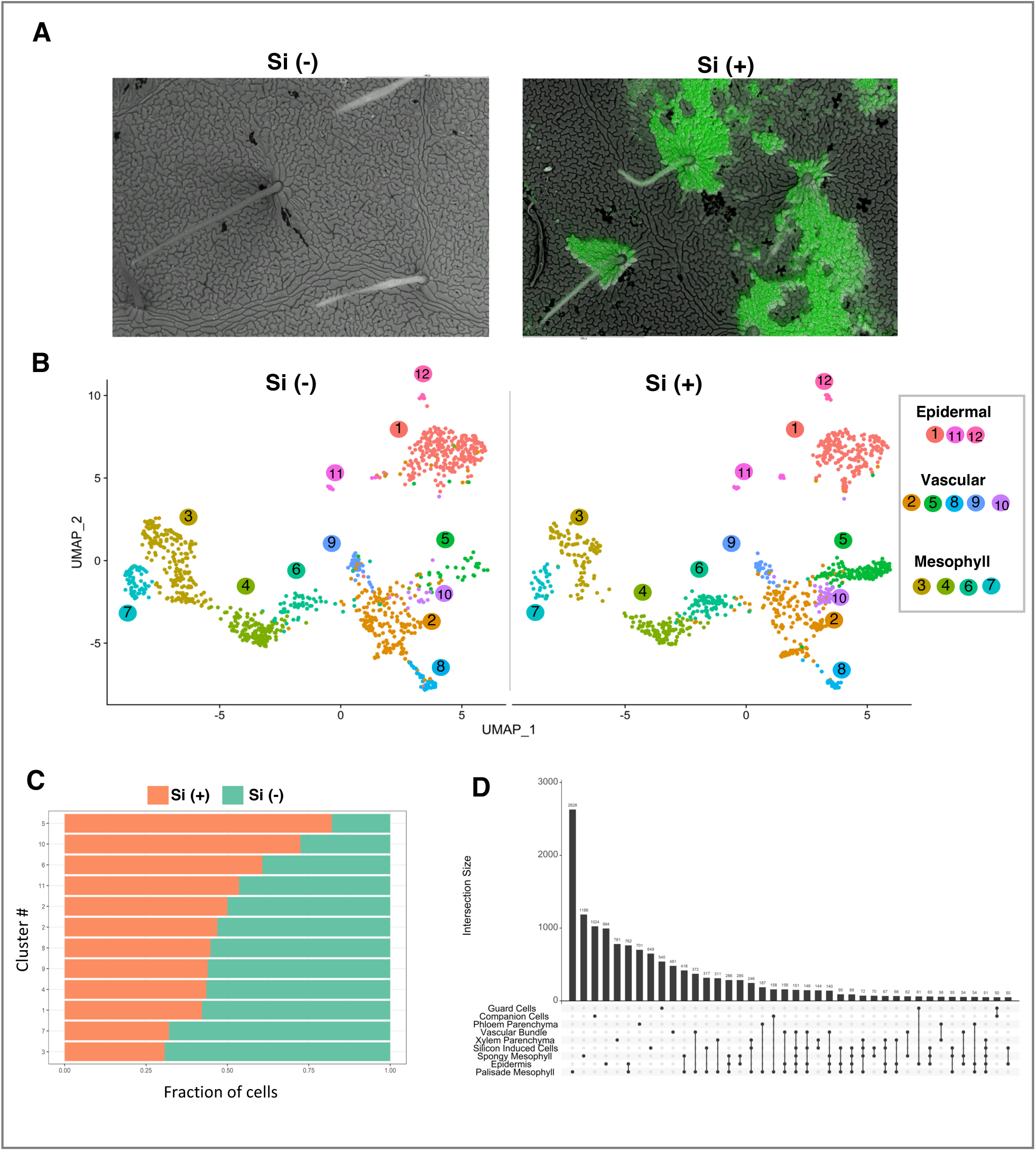
Single nucleus RNA-sequencing (SnRNA-seq) profiles of soybean leaves treated with Silicon. **(A)** Images obtained through scanning electron microscopy combined with EDS spectroscopy. Green color depicts Silicon (Si) accumulation in leaf and trichomes. (**B)** Color-coded visualization of Si treated (Si+) and untreated (Si-) soybean leaf tissue nuclei by Uniform Manifold Approximation and Projection (UMAP) plot based on transcriptomic profiles. These nuclei were categorized into 12 distinct clusters. (**C)** Fractions of nuclei in Si+ and Si-leaf samples. **(D)** Upset plot of expressed genes in different cell clusters in soybean leaves. The left bar plot illustrates the number of enriched DEGs for each cell type, while the top bar plot displays the count of enriched DEGs.

### Identification of “Silicon-induced” cell cluster

To identify preliminary cell populations, Arabidopsis orthologs of cluster-specific genes (Procko et al., 2022; Tenorio Berrío et al., 2022) in conjunction with Gene Ontology (GO) term enrichment analysis was applied (Figure 2A, 2B, 2C, and Supplemental Table S1A). Of the 12 clusters identified (Figure 1B, Supplementary Fig. S3B), nine over-arching cell types were identified. Epidermal cells were clustered into three sub-clusters (#1, 11, and 12-indicated on the figure 2) and exhibited known pathways related to wax and cuticle-formation, some of them include *GmABCG32*, *GmCER4*, *GmKCS10*, *GmGPAT8*. Cluster #12 represented guard cells and genes involved in stomatal differentiation. Through the exclusive expression of genes found within clusters #2, #5, #8, #9, and #10, we identified these cells as “vascular cells” based on our analysis. Specifically, within this group, clusters #8 and #9 were identified as companion and phloem parenchyma cells, respectively. Vascular cell marker genes involved in phloem loading (*GmSUC2*, *GmHPT1*), sap components (*GmLOX1),* and molecular transportation (*GmACA10&12, GmHIPP3)* processes were identified. Clusters #5 and #10 were annotated as xylem-specific cells. However, the number of cells was disproportionately observed in Si+ treatment (Figure 1B and 1C). Interestingly, cluster #5 was nearly absent in control (Si-) samples and is hereafter referred to as “Si-induced cells.” This designation is supported by the well-established phenomenon wherein Si is transported through the xylem before being unloaded into the leaves and thereby, transported to the epidermis (Yamaji et al., 2015). Furthermore, based on the genes involved in photosynthetic and light harvesting complexes (*GmLHCA4*, *GmLHB1B1*, *GmLHCB5*, and *GmLHCB6*) and photosystem I and II genes (*GmPSAK*, *GmPSAE-2*, *GmPSAH2* and *GmPSBY*, *GmPSBX*, and *GmPSBR*), the mesophyll cells were identified. Clusters #3, 4, 6, and 7 were annotated as mesophyll cells (Figure 1B and 2C). Furthermore, these cell types were characterized as palisade mesophyll cells (clusters 3 and 7) and spongy cells (clusters 4 and 6) using Arabidopsis leaf scRNA-seq data (Procko et al., 2022).

**Figure 2.**
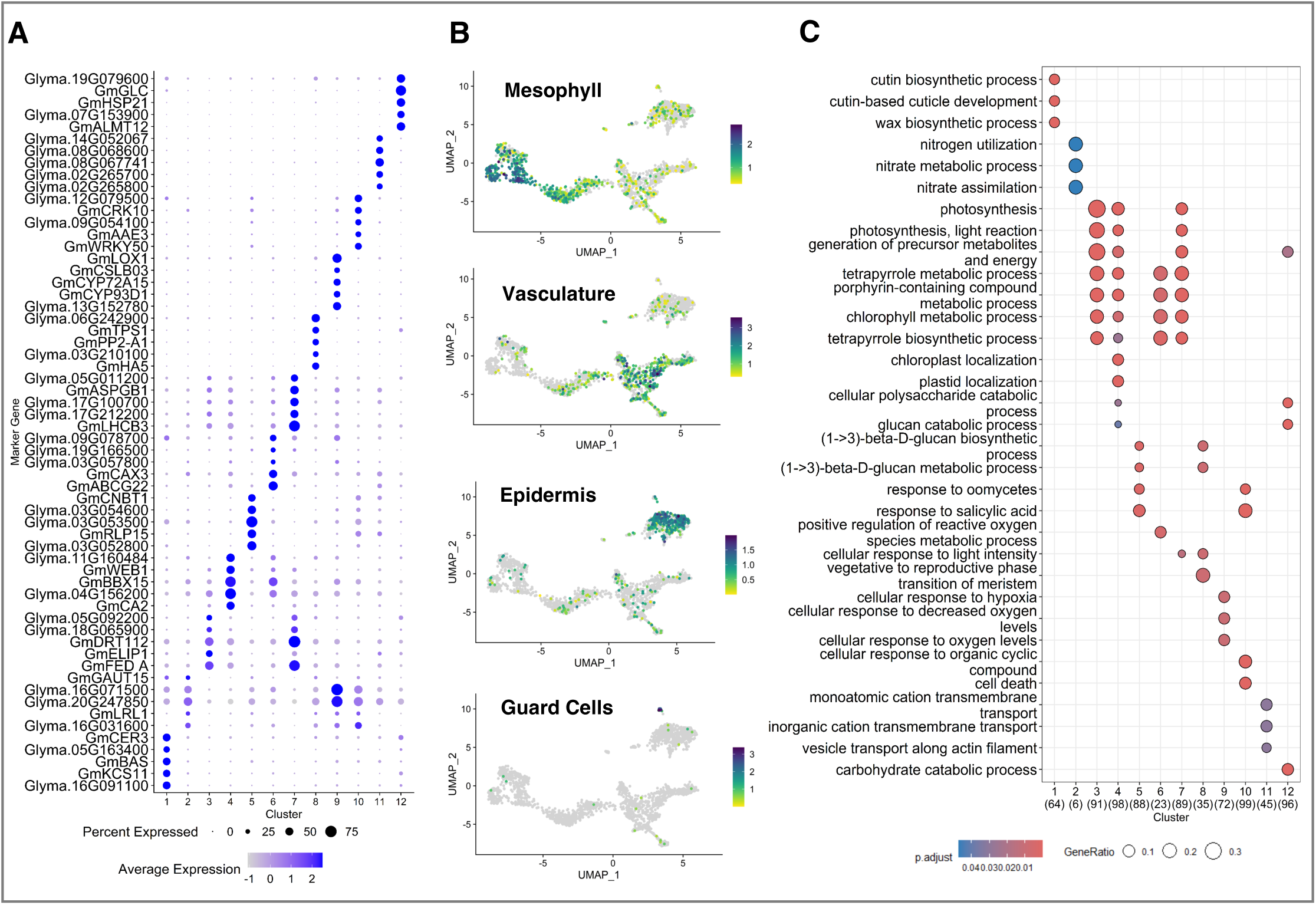
Identification of marker genes for different clusters in soybean leaf tissues. **(A)** Dot plot depicts the top marker genes of distinct clusters. The size of the dots depicts the percentage of cells in each cluster that express the gene. The colors exhibit the relative gene expression. (light blue = lower expression, dark blue = higher expression). A comprehensive list of all marker genes is accessible in Table S1. **(B)** UMAP plot shows detailed expression of selected representative marker genes for overarching cell type clusters such as Mesophyll (*GmFBA2*, *Glyma.04G065600*, *GmPSAH2*, *GmFED-A*, *Glyma.18G057900*), Vascular (*Glyma.16G031600*, *Glyma.15G160300*), Epidermis (*GmMCM4*, *Glyma.16G091100*, *GmCSY3*), and Guard Cells (*GmALMT12*, *Glyma.07G153900, GmHSP21*). The color depicts relative gene expression (dark purple = higher expression, Green = medium expression, Yellow = lower expression). **(C)** Gene Ontology (GO) enrichment analysis for the marker genes within each cluster. X-axis shows GO terms, while Y-axis denotes different cell clusters and numbers in parentheses are the number of genes significantly expressed in each cluster.

### Discerning the cellular responses to Si treatment

Previous studies have investigated the effects of Si treatment on the expression of genes involved in both abiotic and biotic stress tolerance (Kurabachew et al., 2013; Wang et al., 2017). However, the precise mechanism by which Si treatment regulates gene expression across various cell types has remained to be explored. Our study aims to elucidate the specific changes in transcript abundance within individual cell-type clusters induced by Si treatment. Utilizing transcriptome atlases generated from both untreated and Si-treated samples, we identified 4,327 differentially expressed genes (DEGs) across all cell types in response to Si treatment (Supplemental Table S1B).

To discern the impact of Si+ treatment on specific cell types, we conducted a comparative analysis of UMI within our snRNA-seq dataset. This analysis revealed a notable increase in UMI counts for Si-induced xylem cells, phloem parenchyma, and epidermal cells. In contrast, a decrease in UMI values was observed for companion, guard, and mesophyll cells (Figure 3A). We identified DEGs between Si-and Si+ treatments and performed Gene Ontology (GO) and GO term interaction analysis of the DEG sets (Figure. 3B, Supplementary Fig. S4, S5, and S6). The number of DEGs was significantly increased in Si+ treatment wherein vascular cells (Si-induced cells) and epidermal cells were the most affected cell types and support the fact about Si transport from vascular tissues and deposition in epidermal cell layers.

**Figure 3.**
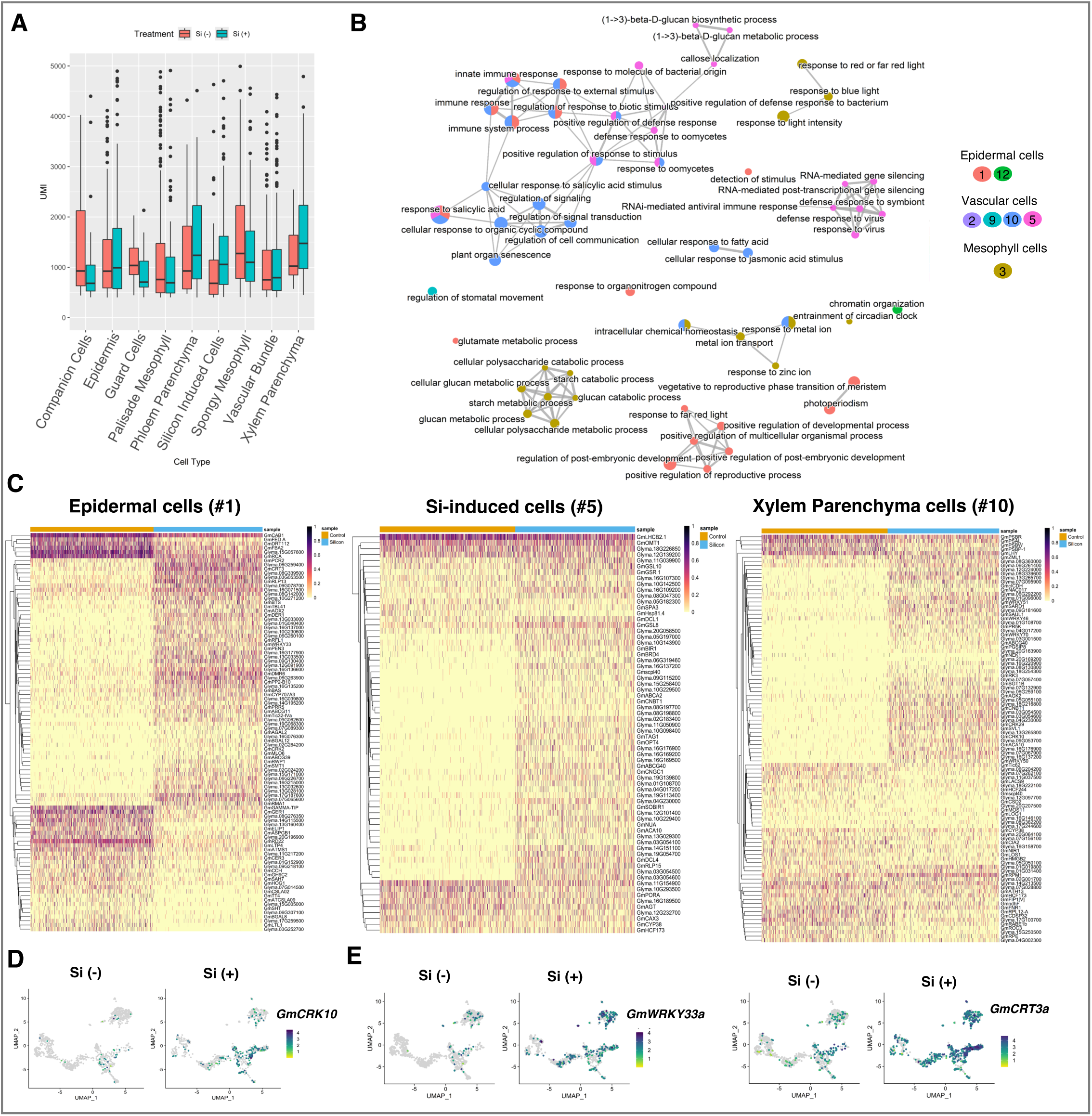
Silicon treatment responses in different cell types. **(A)** Dissemination of total UMI for different cell types in integrated SnRNA-Seq data. **(B)** GO enrichment (biological process) interactions in distinct cell types upon Si (+) treatment in soybean leaf tissues. **(C)** Heatmap depicts differentially upregulated genes in epidermal, Si (-) induced and xylem parenchyma cell clusters after Si (+) treatment. Red color shows upregulation, while yellow color denotes downregulation of gene expression. Expression values were shown in Log_2_FC and P value >0.01. **(D)** UMAP plots display expression of *GmCRK10* at Si (-) and after Si (+) treatment. **(E)** UMAP plot depicts transcript abundance of putative phytoalexin biosynthesis and signaling genes *GmCRT3a* and *GmWRKY33a* upon Si treatment. The color depicts relative gene expression (dark purple = higher expression, Green = medium expression, Yellow = lower expression).

In response to Si treatment, vascular cells, specifically Si-induced cells in cluster #5, exhibited a significant enhancement of GO terms associated with immune response, including NBS-LRR (NLR) pathway genes and WRKY family transcription factors (TFs), as well as defense responses against oomycetes and viruses. Additionally, the GO term “RNAi mediated antiviral response” showed prominent enrichment in these cells (Supplementary Fig. S4). Whereas two other vascular cell types (clusters #10 and #2) displayed enrichment of GO terms related to ‘response to biotic stimulus’, ‘salicylic acid signaling’, and ‘innate immune response’ upon Si+ treatment (Supplementary Fig. S4). Notably, epidermal cells under Si+ treatment exhibited enrichment of GO terms associated with phytoalexin biosynthesis and response to chitin, both known to trigger immune responses (Ahuja et al., 2012; Sánchez-Vallet et al., 2015; Pusztahelyi, 2018) (Supplementary Fig. S5). Overall, GO term interaction analysis revealed potential interactions among genes involved in these processes, suggesting a coordinated response to Si treatment aimed at enhancing biotic stress tolerance in plants (Figure 3B).

To gain further insights into cell-type-specific gene expression under Si treatment, we constructed expression heatmaps for three key cell types: epidermis (cluster 1), xylem parenchyma cells (cluster #10), and Si-induced cells (cluster #5) (Figure 3C). The heatmaps revealed differential upregulation of 97 genes in epidermal cells, 68 genes in Si-induced cells, and 100 genes in xylem cells following Si+ treatment (Figure 3C). A predominant portion of the upregulated genes in xylem parenchyma and epidermal cell clusters upon Si+ treatment were associated with defense responses, immune responses, protein kinases, and NLR pathway genes. Remarkably, the Arabidopsis homolog encoding *Cysteine-rich receptor-like kinase10* (*CRK10*; *Glyma.09G151400*), known for its involvement in *Fusarium oxysporum* resistance (Piovesana et al., 2023), exhibited high expression levels in both epidermal (cluster 1) and Si-induced cells (cluster 5) (Figure 3C and 3D). In Arabidopsis, this plasma membrane-associated protein is expressed in vascular tissues, suggesting its role in plant-pathogen interaction. Similarly, transcription factor (*WRKY33*) and gene (*CALRETICULIN-3a; CRT3a)* associated with phytoalexin biosynthesis were highly expressed in epidermal and vascular cells in Si+ treatment (Figure 3E; Supplementary Fig. S7). Phytoalexins are vital antimicrobial secondary metabolite compounds synthesized in plants in response to biotic stress (Kuć and Rush, 1985; Fawe et al., 1998; Ahuja et al., 2012) Several functional studies have demonstrated the role of *CRT3a* and *WRKY33*a in mediating production of phytoalexins and eventually conferring resistance against fungal diseases (Mao et al., 2011; Matsukawa et al., 2013; Zhou et al., 2020; Ahmed and Kovinich, 2021; Tao et al., 2022).

### Deciphering Si deposition in soybean (dicot) system

The Si uptake, transport, and deposition mechanisms vary among plant species. Si is generally deposited in specialized cells of monocot species such as rice, sorghum, barley, and maize (Kumar et al., 2017; Kumar et al., 2020). Conversely, in dicot species such as Cucurbitaceae (pumpkin, watermelon) (Abe, 2019), sunflower (Van der Ent et al., 2020), soybean (Rasoolizadeh et al., 2018), tomato (Zexer et al., 2023), and broad-leaf trees (Ge et al., 2020), Si is deposited in patches at the trichome base, subsequently spreading to epidermal cell layers. Si deposition in plant cells creating a typical structure of amorphous Si is also called as phytoliths and is used for archaeological tracing (Ge et al., 2020). The reason behind Si deposition in dicots in patches rather than a uniform distribution remains unclear. However, several factors may contribute to this phenomenon, including parallel venation in monocots compared to net-like (reticulate) veins in dicots, which may affect Si unloading in leaves. Additionally, the transpiration rate, dynamics of xylem flow, or preferential unloading of Si in leaves could influence Si deposition. Our observations indicate a significantly higher number of cells in cluster #5 upon Si+ treatment, suggesting that these cells were explicitly differentiated from vascular cells (xylem parenchyma cells) and developed into putative Si-induced cells before unloading Si in the epidermal cells (Figure 4A, 4B and 4C). The UMAP plot of vascular cell types revealed a proximity between xylem cells (#10) and Si-induced cells (#5) (Figure 4C).

**Figure 4.**
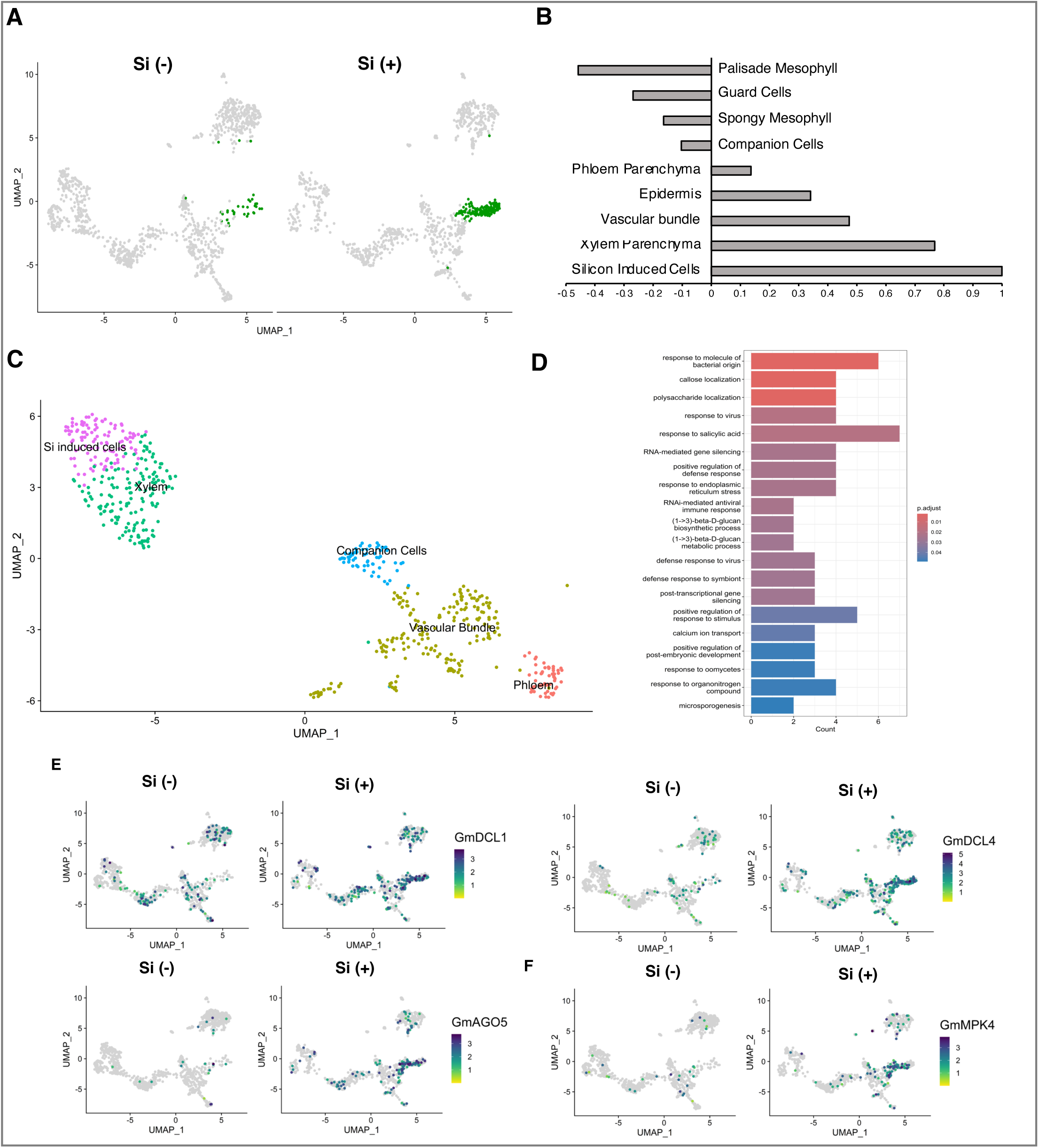
SnRNA-Seq profile of putative silicon-induced cells. **(A)** UMAP plot shows representation of nuclei in Si-induced cell cluster (colored as green) in Si (-) and Si (+) treated soybean leaf samples. Remaining clusters are shown in gray color. **(B)** A bar graph represents the similarity between Si-induced cells and remaining cell type clusters after Si (+) treatment. **(C)** UMAP plot exhibit nuclei clusters in vascular and Si-induced cells. **(D)** GO terms of DEGs in Si-induced cell cluster. E, UMAP shows representative marker genes for individual cell type clusters. **(E)** UMAP plot exhibits expression of RNAi mechanism (PAZ domain) genes (*GmDCL1, GmDCL4, GmAGO5*) in Si (-) and Si (+) treatment. **(F)** UMAP shows expression of *GmMPK4* gene. The color depicts relative gene expression (dark purple = higher expression, Green = medium expression, Yellow = lower expression).

Subsequent gene expression analysis in Si-induced cells revealed that eleven out of twenty enriched GO terms identified in this cluster were directly linked to the defense response against fungi, viruses, and bacteria (Figure 4D, Supplementary Fig. S4 and S8). Additionally, an association map was generated to identify the most distinct genes expressed exclusively in Si-induced cells (Supplementary Fig. S9A). This analysis unveiled several pathogenesis-related genes such as NLRs, RPS3/RMP1, Dicer-Like protein (GmDCL1, GmDCL4), Argonaut (GmAGO5), and mitogen-activated protein kinases (*GmMPK4*) among others that were highly expressed in Si-induced cells upon Si+ treatment (Fig. 4E, Supplementary Fig. S9B). This compelling evidence suggests that Si+ treatment primes plants to activate their host defense mechanisms, enhancing their ability to combat various pathogens.

### Perturbations of defense-responsive genes upon Si treatment

Si treatment significantly affected immune receptor genes such as RLP (Receptor-Like Protein) and NLR (NBS-LRR; nucleotide-binding domain leucine-rich repeat). RLPs function as cell surface receptors responsible for detecting pathogen-associated molecular patterns (PAMPs), thereby initiating a fundamental defense mechanism termed PAMP-triggered immunity (PTI). These receptors play a crucial role in regulating plant defense responses against various pathogens, including fungi, oomycetes, and bacteria (Jamieson et al., 2018). We identified 6 RLP genes that were highly expressed in vascular cells upon Si+ treatment (Figure 5A, 5B and Supplemental Table S1C). Remarkably, the RLP genes such as *GmRLP9*, *GmRLP13* and *GmRLP56* showed higher expression in xylem and Si-induced cells after Si+ treatment (Figure 5C).

**Figure 5.**
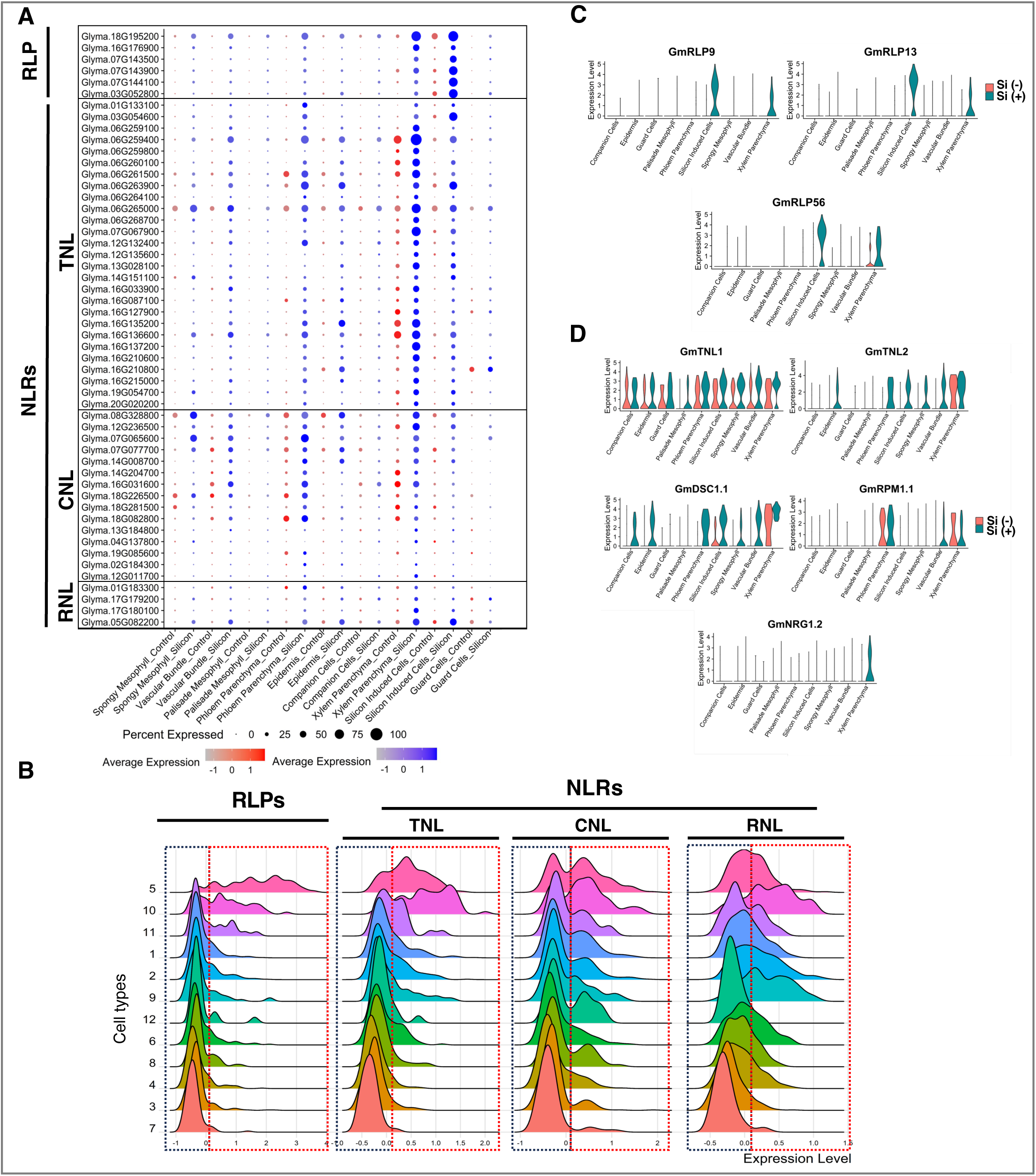
Transcriptional regulation of RLP and NLR genes in different clusters upon Si treatment. **(A)** A dot plot depicts expression of RLP and NLR immune receptor genes in different clusters upon Si (+) treatment. The size of the dots shows the percentage of cells in each cluster that express the gene. The colors exhibit the relative expression. The Red dots indicate control (Si-) cells while blue dots indicate Si+ treated cells. Control is on the left and Silicon is on the right for each tissue-type pair. **(B)** A Violin plot exhibits expression level of RLP and NLR genes in all different cell type clusters. The height of each peak denotes the number of genes, while the width of each peak depicts expression level. The X-axis shows expression level, while the Y-axis denotes different clusters. **(C)** A violin plot illustrates transcript abundance of RLP genes namely *GmRLP9*, *GmRLP13*, and *GmRLP56*. D, Violin plot shows expression level of NLRs namely *GmTNL1*, *GmTNL2*, *GmDSC1.1*, *GmRPM1.1*, *GmNRG1.2*. For C-D, the Si (-) and Si (+) samples are shown side-by-side for each tissue type. Treatments are differentiated by violin color.

Similarly, NLRs are intracellular immune receptors that perceives cytoplasmic effectors secreted by pathogens and initiate effector-triggered immunity (ETI) (Cui et al., 2015; Ngou et al., 2022). NLRs are categorized into three different classes based on their diverse N-terminal domains, namely, toll/interleukin-1 receptor/resistance protein NLR (TIR-NLR/TNL), coiled-coil-NLR (CNL/CC-NLR) and RPW8-like helper NLR (RNL) (Kourelis et al., 2021). In the soybean genome, 319 genes were identified as potential NLR genes (Kang et al., 2012). In this study, we identified 46 NLR genes (27 TNLs, 15 CNLs and 4 RNLs), that were highly expressed in Si+ treated plants and confined mainly to epidermal and vascular cell clusters (#1, 2, 5, 9, 10, and 11) (Figure 5A, 5B and Supplemental Table S1C). In several plant species, including soybean and Arabidopsis, it has been shown that the NLRs genes such as *Rpp1*, *RPS11*, *DSC1, RPM1,* and *NRG1.2* are involved in fungal, oomycetes, bacterial, and viral defense (Boyes et al., 1998; Peart et al., 2005; Castel et al., 2019; Pedley et al., 2019; Wang et al., 2021). Moreover, in soybean the TNL class gene *GmTNL2* was identified as a candidate gene for resistance to Asian soybean rust (*P. pachyrihizi*) (Yu et al., 2015). In our snRNA-seq dataset, the TNL class genes such as *GmTNL1*, *GmTNL2* and *GmDSC1.1* exhibited higher expression in epidermal, vascular and Si-induced cells upon Si+ treatment (Figure 5D), whereas in the case of CNL and RNL classes *GmRPM1.1* and *GmNRG1.2* exhibited higher expression specifically in xylem and phloem parenchyma cells (Figure 5D). These results indicate that RLP and NLR immune receptor pathways were induced upon Si+ treatment, mainly in vascular cells in soybean leaf.

Salicylic acid (SA) is a signal molecule (plant hormone) involved in systemic acquired resistance (SAR) by triggering the plant immune system and activating defense and pathogenesis-related genes (Stroud et al., 2022; Rossi et al., 2023). Previously, researchers have used various formulations of Si and SA to improve abiotic stresses by inducing antioxidant enzymes (Khan et al., 2019; Yang et al., 2021; Shohani et al., 2023). However, whether Si treatment can induce SA biosynthesis or its upstream signaling genes in plants is elusive. In our dataset, we observed cell-specific expression of SA biosynthesis and signaling genes (Supplementary Fig. S10A). SA biosynthesis occurs through isochorismate (ICS) and the phenylalanine (PAL) pathway (Klämbt, 1962; Wildermuth et al., 2001; Huang et al., 2020). In soybeans, both PAL and ICS pathways contribute equally to SA biosynthesis (Shine et al., 2016). However, we did not observe the induction of ICS and PAL pathway genes upon Si+ treatment, while these genes were expressed only in control (Si-) treatment (Supplementary Fig. S10A and S10B). Similarly, the MES (methyl esterase) gene, involved in converting inactive Methyl-SA to Active SA (Vlot et al., 2008), was expressed in mesophyll cells of both Si-and Si+ treatment. But (interestingly), *GmSARD1,* a regulator for *ICS1* in SA synthesis, was highly expressed in vascular cells (#10). This observation suggests that although Si+ treatment does not induce the core SA pathway genes, the upstream master regulators of the SAR pathway, including *GmSARD1*, *GmPAD4*, and *GmWRKY40*, were highly induced in vascular and guard cells upon Si+ treatment. Expression of these genes has been reported to increase pathogen resistance, stomatal closure, and SAR through SA biosynthesis in plants (Zeng et al., 2010; Prodhan et al., 2018). Furthermore, genes involved in SAR downstream to SA biosynthesis pathways such as *GmNPR3* (Non-expression of PR protein), *GmTGAs* (TGACG cis-element binding), and *GmWRKY70* were expressed in vascular and epidermal cells, whereas *GmPR1s* (*Pathogenesis related-1*) were expressed in vascular and mesophyll cell types upon Si+ treatment.

### GmHiSil2c regulates Si unloading in epidermal cell layers

Silicon transporters have been well-characterized in monocots, mostly in rice. Si transport is mediated by two types of transporters, *Lsi1* (influx) and *Lsi2* (efflux). Lsi1 facilitates the translocation of Si from the soil into root cells via a gradient mechanism, functioning as a passive transporter. Conversely, *Lsi2* acts as an active transporter, facilitating the transport of Si from root cells to the vascular stream (Ma et al., 2006; Mitani et al., 2008; Ma, 2010; Ma and Yamaji, 2015; Deshmukh and Bélanger, 2016; Coskun et al., 2021; Mitani-Ueno et al., 2023). In a recent genome wide association study (GWAS) (Dhingra et al., 2024), we identified that Si accumulation in soybean leaves is controlled by a major QTL on Chromosome (Chr) 16, which harbors three efflux Si transporter genes (*GmHiSil-2a*, -*2b* and *-2c*). To understand cell-specific expression of these genes identified on Chr. 16 and other putative Si influx aquaporin family transporters (Deshmukh et al., 2013), the snRNA-seq data was explored.

Among the several putative Si transporters in soybean (Supplemental Table S1D), the two influx Si transporters *GmNIP2-1* (*Lsi1*), *GmNIP2-2* (*Lsi1*) were found to be expressed in our datasets (Figure 6A). On the other hand, among the three Si efflux transporters identified in GWAS (Dhingra et al., 2024), only *GmHiSil2c* (*Lsi2*) was highly expressed in our snRNA-seq dataset. The UMAP plot showed that the expression of *GmNIP2-1* and *GmNIP2-2* was notably higher in Si-(control) leaves, particularly in the epidermal cells, whereas their expression decreased following Si+ treatment (Figure 6A). Meanwhile, *GmHiSil2c* was exclusively expressed in leaf epidermal cells upon Si+ treatment. Additionally, the expression has been detected to a lesser extent in vascular cells (Figure 6A). Furthermore, in an independent experiment we validated the expression of these genes using qRT-PCR analysis and their expression pattern was as similar as snRNA-seq data (Figure 6B). These findings suggest that *GmHiSil2c* acts as a functional Si transporter specific to leaves, possibly playing a role in Si loading into the space between the epidermis and cuticle bilayer. Subsequently, to determine the subcellular localization of these Si transporters, the coding sequences (CDS) of *GmNIP2-1*, *GmNIP2-2*, and *GmHiSil2c* were fused with the green fluorescent protein (GFP) at the C-terminal end and expressed under the control of constitutive promoter (CmYLCV). Transfection (PEG-mediated) of these transporters was performed in *N. benthamiana* protoplast. Subsequently, confocal microscopy analysis revealed that these transporter-GFP fusions were localized to the cell membrane, whereas the control construct (CmYLCV::GFP) was expressed and localized to the cell membrane, nucleus, and cytoplasm (Figure 6C).

**Figure 6.**
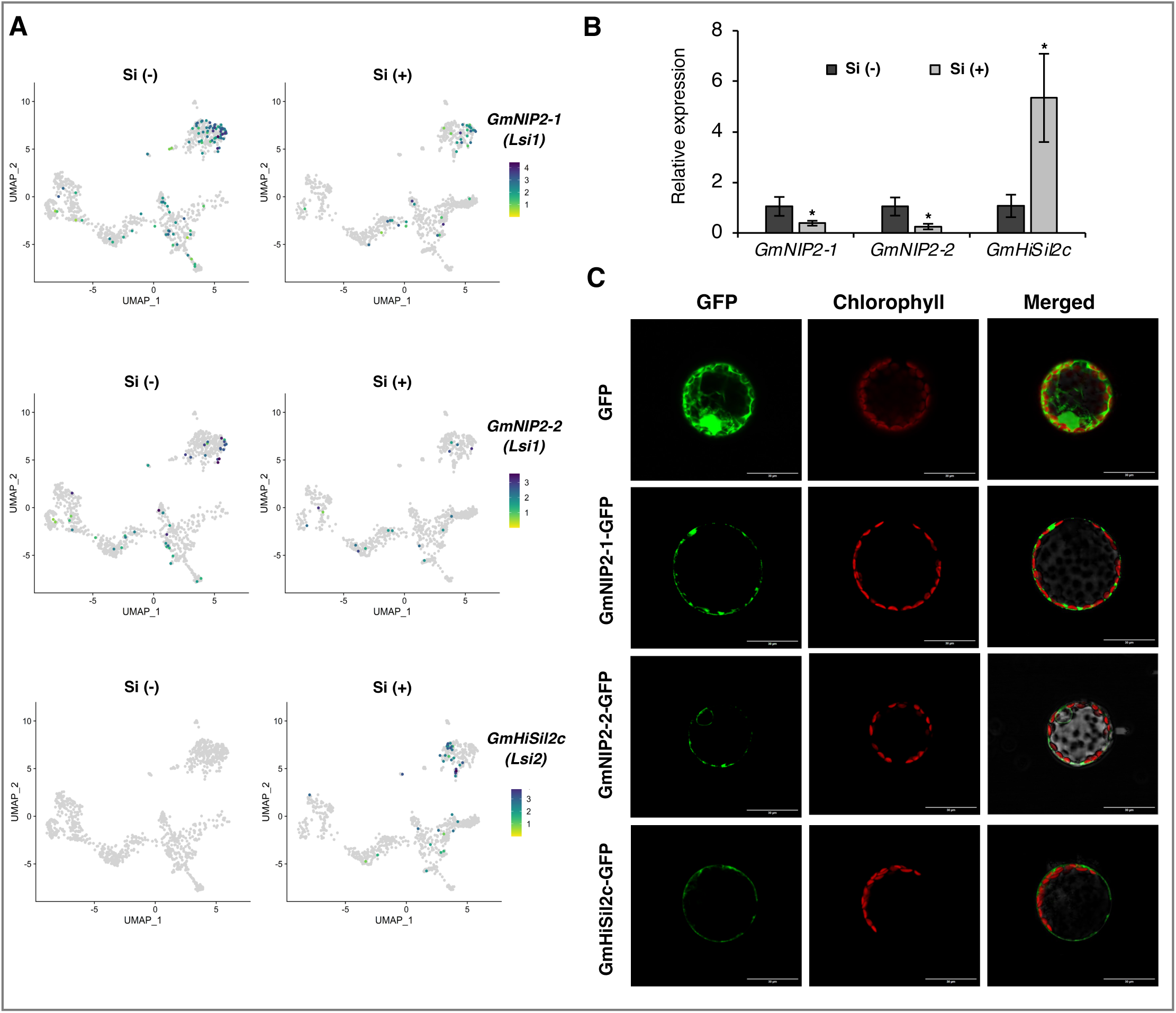
Identification of expression pattern of Si transporters in different cell types. **(A)** UMAP plot exhibits expression of Si transporters *GmNIP2-1* (Upper plots), *GmNIP2-2* (Middle plot), *GmHiSil2c* (Lower plots) in all clusters in leaves of Si (-) and Si (+) treated plants. **(B)** The qRT-PCR expression of *GmNIP2-1*, *GmNIP2-2*, and *GmHiSil2c* in leaves of Si (-) and Si (+) treated plants in hydroponic system. Expression values are shown in fold change. Gene expression is relative to reference gene *GmElf1b*. The data shows mean for three biological replicates ± SD. Asterisks denotes a significant difference between Si (-) and Si (+) treatment (Students t-test; *, P < 0.05). **(C)** Subcellular localization of Si transporters GmNIP2-1-GFP; GmNIP2-2-GFP; GmHiSil2c-GFP; and only GFP (as control) in protoplasts of tobacco leaves. *CmYLCV* was used as constitutive promoter for only GFP and Si transporters plus GFP fusion.

## Discussion

### Understanding cellular heterogeneity upon Si accumulation in leaves

In plant biology, numerous reports substantiate the importance of mineral nutrients on plant development, disease resistance, abiotic stress tolerance, seed quality, and net productivity (Dhingra et al., 2024). The availability of mineral nutrients to plants is a dynamic and complex process affected by various factors. These factors encompass the inherent gene pool governing nutrient uptake and translocation, soil composition, physical and chemical properties, and other environmental variables. Among several major and minor nutrients, the role of Si in plants is often overlooked. Although, Si is not classified as an essential plant nutrient, it plays a beneficial role in plant development (Pavlovic et al., 2021), abiotic stress tolerance (Gao et al., 2005; Hamayun et al., 2010; Wang et al., 2021), and biotic stress alleviation (Fauteux et al., 2005; Fauteux et al., 2006; Rasoolizadeh et al., 2018; Thakral et al., 2021). The molecular mechanisms governing Si intake and translocation have been extensively examined in rice (Ma et al., 2006; Mitani et al., 2008; Ma, 2010; Mitani-Ueno et al., 2023). However, its role in various dicot crop plants including soybean is not well elucidated. In recent studies, Dhingra et al. (2024) and Belanger et al. (2019), a major QTL for Si uptake in soybean leaves has been identified. However, there is a lack of a comprehensive molecular understanding of Si uptake and translocation in dicot species, including soybean. Also, previous studies in other plants primarily relied on bulk transcriptomics, a methodology inherently unable to discern cellular heterogeneity and prone to overlooking cell-specific gene expression (Fauteux et al., 2006; Chain et al., 2009; Ma et al., 2016; Zhu et al., 2019; Jiang et al., 2022).

Single-nuclei-based transcriptomics is a powerful technology for elucidating the dynamics of biological processes and understanding the diversity among distinct cell types. The snRNA-seq techniques have enabled researchers to analyze complex traits at a single-cell resolution, particularly emphasizing tissue-specific single-cell atlas, tissue developmental processes, plant-microbe interactions, and abiotic stresses (Kim et al., 2021; Cervantes-Pérez et al., 2022; Tenorio Berrío et al., 2022; Li et al., 2023; Nobori et al., 2023; Sun et al., 2023; Tang et al., 2023). To comprehensively elucidate Si deposition in soybean leaves and its consequential effects on genes specific to individual cells and their associated gene networks, we employed snRNA-seq technology. Our approach allows us to gain comprehensive insights into the intricacies of the Si mediated biological process in plants. The foundational step in single-cell genomics is the classification of cell types. Remarkably, the identification of leaf-specific cell types in soybean has remained unexplored to date, and to address this critical gap, we undertook a comprehensive approach, leveraging publicly available datasets (Kim et al., 2021; Procko et al., 2022; Tenorio Berrío et al., 2022) and iterative tools such as the R package ClusterFoldSimilarity (González-Velasco et al., 2022) and cross-species integration to classify the diverse cell types within soybean leaves systematically. Previous studies in rice, Arabidopsis, and Maize identified 10 to18 cell types based on different cell type markers (Marand et al., 2021; Wang et al., 2021). In this study, we developed snRNA-seq transcriptome of soybean plants treated with and without Si and successfully unveiled three major tissue types (vascular, epidermal, and mesophyll) that were further subclustered into 12 distinct cell populations (Figure 1). Interestingly, a novel cell cluster #5 termed ‘Si-induced cells’ was identified only in Si+ treated soybean plants, suggesting a unique mechanism of Si deposition in soybean leaves. These unique clusters enabled us to detect cell-specific transcript abundance during Si+ treatment.

### Si treatment unlocks key role of vascular cells in plant defense

After conducting an in-depth characterization of distinct cell types, we identified a complex, yet compelling gene expression pattern was identified (Figures 1, 2, and 3). Importantly, we identified a subset of genes and their GO terms linked to ‘defense response’, ‘plant immunity,’ and response to salicylic acid and phytoalexin (Figure 3). There are several mechanisms through which Si ameliorates biotic stress in plants, such as serving as a physical barrier or apoplastic obstruction, enhancing defense-responsive metabolites, hormones, and antimicrobial compounds (Phytoalexins), and activating defense-responsive molecular cascades (Fauteux et al., 2006; Wang et al., 2017). Interestingly, some studies also suggested that Si alone has no impact on plants’ metabolism in a controlled, unstressed condition (Cai et al., 2008; Coskun et al., 2019). Our study confirms that Si gets deposited in epidermal cell layers of soybean leaves, creating an apoplastic barrier, but we also observed that Si treatment alone can change the leaf transcriptome and primes the plant to induce defense-responsive genes even in the absence of pathogenic stress.

Consistent with previous research conducted by Brunings et al. (2009), Hao et al. (2021), and Hu et al. (2018), which illustrate that Si supplementation triggers distinct regulatory patterns in the transcriptome of plants in the absence of a pathogen or abiotic stressor. Notably, Hao et al. (2021) reported that Si treatment in wheat elicits the upregulation of numerous genes, particularly highlighting the differential expression of MYB transcription factors. Hu et al. (2018) also provided evidence of Si-induced reduction in phosphorus (P) uptake through the downregulation of P transporter genes, independent of any biotic or abiotic stressors. Nevertheless, it cannot be ruled out that the priming effect through transcriptome could be temporal. Thus, further investigation is warranted to elucidate the implications of these transcriptome changes on downstream outcomes such as protein and metabolite production.

Intriguingly, our study suggested cell-specific expression of defense responsive genes, significantly confined to vascular, particularly in Si-induced cells (cluster #5) upon Si treatment (Figure 4). This observation represents a significant advancement beyond prior knowledge, emphasizing the distinct and specialized role of Si treatment in coordinating molecular responses related to biotic stress and phytohormonal signaling. Particularly noteworthy was the involvement of vascular cells as critical mediators in these intricate regulatory processes initiated by Si treatment (Figure 3C). Although, the precise mechanisms underlying the notable increase in cell abundance within cluster #5 following Si+ treatment remains to be studied (Figure 4A), examination based on cell-type-specific markers revealed that this cluster was closely associated with vascular cells (Figure 4B). Interestingly, it has been reported that both high Si-accumulating (rice and maize) and low Si-accumulating (onion) plants develop new Casparian band cells in roots upon Si treatment (Fleck et al., 2015). This observation suggests that in dicot, pre-existing vascular cells may undergo differentiation, giving rise to new cell types (cluster #5, Si-induced cells). However, this hypothesis necessitates dedicated experimentation to validate and further explore the underlying mechanisms. Moreover, this Si-induced cluster exhibited a unique gene expression pattern significantly enriched in genes related to plant-pathogen interaction. The higher transcript abundance of genes associated with RNAi silencing machinery, defense response and certain immune receptors such as RLP and NLR genes were specific to this cluster. It is well-studied that RNAi silencing machinery gets activated against pathogen attack such as viral infections and degrade the viral RNA via small interfering RNAs (siRNA), preventing the spread of the infection (Deleris et al., 2006; Mukherjee et al., 2013; Qin et al., 2017). This mechanism is largely conserved in plants and functions through RNA-induced silencing complexes (RISCs), DICER-LIKEs (DCLs), and Argonaute (AGO) proteins.

We identified a significantly higher transcript abundance of RNA silencing machinery genes, such as *GmDCL1*, *GmDCL4*, *GmAGO5* and *GmMPK4*, in Si-induced and other cell types (Figure 4E). Among the prominent biological processes, genes involved in the phytoalexin biosynthesis process were also significantly enriched in epidermal (cluster #1, 11), vascular (cluster #2, 5, 8, 9 and 10) and guard cells (cluster #12) after Si+ treatment. Phytoalexins have antimicrobial properties and are synthesized in response to the onset of stress cues in plants and are involved in pathogen defense response (Fawe et al., 1998; Ahuja et al., 2012). In soybean, glyceollin is a major phytoalexin, underlying isoflavonoid group, and has a crucial role in providing defense against Phytoph*thora sp.* (Schmidt et al., 1992; Fawe et al., 1998; Rodrigues et al., 2004; Cheng et al., 2015). However, the specific molecular pathways occurring in distinct cell types that lead to the synthesis of phytoalexins upon Si treatment remain elusive. In our investigation, we observed the expression of phytoalexin activator genes (*GmWRKY33a, GmWRKY33b, GmWRKY33c*) in epidermal, vascular, and guard cells following Si+ treatment, suggesting their potential involvement in the Si+ primed resistance mechanism. This finding lays the groundwork for further exploration into the underlying mechanisms.

We also identified complementary pathways involving extra-cellular and intra-cellular receptors, such as RLPs and NLRs, marked by a notable upregulation in Si+ treatment (Figure 5A and 5B). RLPs are cell surface receptors involved in the detection of extracellular pathogen molecular signatures, through which they promote pathogen defense response (Bjornson et al., 2021). Specifically, we detected higher expression of *GmRLP56*, *GmRLP13*, and *GmRLP9* genes confined to xylem and Si-induced cells after Si+ treatment (Figure 5C). Similarly, NLR genes, extensively studied in Arabidopsis and various plant species, play a pivotal role in recognizing and responding to specific pathogen effectors. These genes initiate defense mechanisms through hypersensitive response (HR) and SAR (DeYoung and Innes, 2006). A recent study (Tang et al., 2023) applied single-cell transcriptomics to reveal cell-type specific responses to fungal infection in Arabidopsis leaves. Notably, in agreement of our study, they reported the enrichment of intracellular immune receptors (NLRs) in vascular cells (Tang et al., 2023). In our study, snRNA-seq provided a subtle perspective, enabling the detection of distinct expression patterns of NLR immune response genes based on their classification *GmTNL1, GmTLN2, GmDSC1.1* (TNLs), *GmRPM1.1* (CNL) and *GmNRG1.1* (RNL) in cell-types specific manner upon Si+ treatment (Figure 5D).

Salicylic acid (SA), a central player in plant innate immunity, orchestrates various defense responses against biotic attacks. SA acts locally at the infection site and systemically throughout the plant, commencing resistance, hypersensitive responses, and cell death to cease pathogen spread (Ding and Ding, 2020). While the influence of Si on SA has not been extensively investigated in previous studies, our observations reveal a subtle alteration in the expression of SA signaling genes post Si+ treatment, without significant changes in the core SA biosynthesis genes (Supplementary Fig. S10). The upregulation of SA signaling genes (*GmPAD4, GmSARD1,* and *GmWRKY40*) in vascular cells following Si+ treatment supports the notion of priming the SA pathway. However, Si induces the expression of core SA biosynthesis genes (*GmICS* and *GmPAL*), suggesting an unknown negative regulator of SA. This finding is consistent with previous studies by Rasoolizadeh et al. (2018) and (Ye et al., 2013) which reported that Si+ treatment alone did not induce endogenous SA biosynthesis genes in soybean roots and jasmonate (JA) genes (OsAOS and OsCOI1) in rice leaves. Furthermore, our findings support that while pathogen infection is necessary to induce the expression of core SA and JA biosynthesis genes and their accumulation in plant cells, we also conclude that Si+ treatment alone may still induce the expression of SA signaling and receptor genes, such as *GmSARD1*, *GmPAD4*, *GmWRKY40*, *GmNPR*, *GmPR* and *GmTGAs* (Supplementary Fig. S10). In summary, pathogens employ diverse strategies to infiltrate host plant tissues, establishing robust colonization across various cell types. Particularly, vascular tissues are the primary destination for a variety of pathogens (fungi, oomycetes, bacteria etc.) seeking their nutrients. Our finding further substantiates the vital role of Si absorption and modulation of gene expression networks in vascular cells to induce and prepare defense response in plants.

### Unveiling Si deposition in soybean leaves

Silicon uptake in plants occurs primarily through the root epidermal cells, facilitated by specific transporters. The influx transporter *Lsi1*, a member of the aquaporin family, mediates Si uptake from the soil, while the efflux transporter *Lsi2*, belonging to the anion transport protein superfamily, facilitates Si efflux from the root cells into the xylem vessels. These transporters have been extensively characterized in monocot species such as rice and maize (Ma, 2010). To understand the expression patterns of putative Si influx and efflux transporters in soybean (Deshmukh et al., 2013; Deshmukh and Bélanger, 2016), the snRNA-seq data generated in this study was explored in-depth. Among all putative Si transporters investigated, only two influx transporters, *GmNIP2-1* and *GmNIP2-2* (homologous to *OsLsi1*), and one efflux transporter, *GmHiSil2c* (homologous to *OsSiet4*), were expressed in soybean leaf snRNA-seq datasets. Furthermore, the expression of these three genes was validated using qRT-PCR study, revealing relatively higher expression of *GmNIP2-1* and *GmNIP2-2* in epidermal and vascular cells of control (Si-) leaves compared to those subjected to Si+ treatment (Figure 6A). Sub-cellular localization of these genes further confirmed their role as transmembrane proteins. The reduction in expression of *Lsi1* influx transporters (*GmNIP2-1* and *GmNIP2-2*) following Si+ treatment suggests a potential gradient-based transport mechanism, similar to observations in rice (Mitani et al., 2008). Interestingly, in rice, Si+ treatment downregulates the expression of both *Lsi1* and *Lsi2* genes (Mitani et al., 2008; Mitani-Ueno et al., 2016). However, our findings in soybeans revealed that *GmHiSil2c* was highly expressed in Si+ treatment only in epidermal and vascular cells (Figure 6A). These results suggest that while some aspects of Si transport may be conserved across plant species, there are notable differences in the regulatory responses to Si+ treatment between monocots and dicots. Recently, the new leaf-specific Si efflux (*OsSiet4*) has been identified to regulate Si distribution in rice. The knockout mutant of *ossiet4* exhibited a severe wilting phenotype in the presence of Si+ due to disturbed and abnormal Si accumulation in mesophyll cells (Mitani-Ueno et al., 2023). Although, the mechanism of Si efflux transporters has not been studied before in dicot, we identified that *GmHiSil2c* (homolog of rice *OsSiet4*) is mainly expressed upon Si+ treatment (no expression in control (Si-)) in soybean epidermal cells (cluster #1, #11) and its protein localized to the cell membrane (Figure 6C), suggesting the precise distribution of Si in leaf tissues. Overall, these findings contribute to our understanding of Si transport mechanisms in soybean and highlight the potential regulatory mechanisms governing Si uptake and translocation in dicotyledonous plants. Further studies elucidating the regulatory networks controlling Si transporter expression and activity will enhance our ability to manipulate Si uptake and utilization in crops for improved agronomic performance and stress tolerance.

## Materials and Methods

### Plant material, growth conditions, hydroponic system, and Silicon treatment

Soybean cultivar *Hikmok-sorip* (PI 372415) was used in this study. Seeds were grown on germination paper for duration of 5 days; subsequently, seedlings were transplanted onto hydroponic trays containing a solution with half-strength Hogland media at a pH of ∼5.7. The hydroponic system was continuously aerated using an air pump to prevent oxygen deficiency. The hydroponic setup was established in a growth chamber with a photoperiod of 16/8 light/dark, providing 350 PAR light, relative humidity of 65%, day/night temperature regime at 26/24 °C. The nutrient solution in the hydroponic system was replenished once in five days. After 2 weeks of seedling growth, silicon treatment was performed by applying 1.7 mM of K_2_SiO_4_ (Silicon treatment; Si+) for 7 days, while 1.7 mM of KCl (control, Si-) was used as a potassium control (Rasoolizadeh et al., 2020).

### Silicon estimation by SEM-EDS

Silicon content analysis in soybean leaves was conducted through Scanning Electron Microscopy-Energy Dispersive X-ray Spectroscopy (SEM-EDS) as described by Dhingra et al. (2024). The analysis was conducted at an accelerating voltage of 20 kV and 140 Pa under the variable pressure mode using a Hitachi S-3400 microscope equipped with an Oxford EDS system.

### Silicon estimation by XRF

The plant tissues were subjected to a week-long drying process at 45°C, after which they were finely pulverized using the Tissue Lyser II (Qiagen, Cat. No. / ID: 85300). Subsequently, ∼300 mg pellets were prepared from the dried and finely ground samples using a 15T hydraulic press with a 13 mm diameter (Athena Technology, India). Portable X-ray fluorescence (pXRF) analyzer (Bantam Axon, Olympus) was used to quantify mineral element accumulation in each sample as previously described (Otaka et al., 2014). Briefly, the pXRF was set on the analysis mode (Vegetation Analysis) specifically optimized for plant samples by the manufacturer (Bantam Axon, Olympus). The NIST (National Institute of Standards and Technology) samples, including Apple Leaves (NIST1515), Peach Leaves (NIST1547), and Pine Needles (NIST-1575) with known mineral concentrations, were used to generate user factors for each element and as standard control to check the reproducibility of the XRF analysis.

### Nuclei extraction, SnRNA-seq library preparation, and sequencing

Nuclei isolation was performed from Si+ and Si-soybean leaf tissues as previously described (D’Agostino et al., 2023). Briefly, the tissue was ground in a nuclei isolation buffer and incubated briefly on a slow shaker. After incubation the nuclei suspension is passed through 100 µm, 70 µm, 30 µm, and 10 µm cell strainers. The suspension was spun at 1000g for 5 minutes. The nuclei pellet was resuspended in wash buffer and spun again at 1000g for 2 minutes. The nuclei were stained with propidium iodide and counted with the Luna-FL (Logos Biosystems). The nuclei were then loaded onto the 10x Genomics chip as per 10x genomics recommendations. Library construction for Illumina sequencing was performed with the Chromium™ Single Cell 3’ Library & Gel Bead Kit v3.1 protocol (10x Genomics). The single-indexed sequencing and paired-end libraries was carried out on an Illumina™ NovaSeq 6000 platform according to the 10x Genomics references.

### SnRNA-seq data pre-processing, integration, and clustering

Sequencing data was processed through cellranger (v7.2.0) to obtain cell feature counts for analysis. The data was aligned to the Glycine max Wm82.a4. v1 genome. Count data was further analyzed within the R package Seurat v4 (Hao et al., 2021). Nuclei were filtered according to the following criteria: RNA gene count >200, UMI count >400 and <10000, mitochondrial reads >5% per cell, and chloroplast reads <30% per cell. Doublets were removed with scDblFinder (https://www.ncbi.nlm.nih.gov/pmc/articles/PMC9204188/). The data was then normalized by SCTransform and the samples were integrated using 3000 features to reduce batch effect and identify conserved cell types. Dimension reduction of the integrated data was performed by PCA with RunPCA and the UMAP plots were generated using PCs 1 to 40. Leiden clustering was applied (resolution 0.8) to identify the potential cell types. Ubiquitous/nondescript clusters were removed manually and the UMAP and clustering were re-run according to previous specifications. After filtering 2080 total cells were obtained (1104 from control sample and 976 from silicon treated sample) with an average UMI of 1546 and a number of genes of 1005.

### Annotation of Cell types by utilizing ortholog gene markers and Gene Ontology (GO) analysis

Annotation of cell clusters was performed through orthologous expression of Arabidopsis genes and GO term analysis. Markers for each cluster were generated with “FindAllMarkers” at the following specifications: min.pct = 0.25, logfc.threshold = 0.25, min.diff.pct = 0.15. Arabidopsis orthologs were then compared to (Kim et al., 2021) and Leaf SC atlas, VIB Ghent, Belgium (Tenorio Berrío et al., 2022) (https://www.psb.ugent.be/sc-leaf-yieldlab/) to identify cell types. ClusterProfiler (Yu et al., 2011) was used to perform GO term enrichment analysis with default parameter.

### Differential gene expression and Gene Ontology analysis

Differentially expressed genes between control and silicon treatments were found with FindMarkers with the following parameters: min.pct = 0.25, logfc.threshold = 0.25, min.diff.pct = 0.15, group.by = sample, subset.ident = cluster #. Gene Ontology analysis for treatment induced differentially enriched GO terms was performed with clusterProfiler as previously described.

### RNA extraction and gene expression analysis

Total RNA extraction and purification from soybean leaves and roots were carried out using Trizol reagent, followed by the utilization of the Pure Link RNA Mini Kit (Invitrogen). Subsequently, cDNA synthesis and qRT-PCR procedures were conducted following previously established protocols (Le et al., 2012; Devkar et al., 2020). The PCRs were executed using the C1000 Touch PCR with the CFX96 Real-Time Optics Module qPCR System (BIO-RAD, USA). The qPCR primers were designed using Quant prime online tool. *GmElf1b* (Glyma02g44460) served as the internal control for gene expression analysis. All the qPCR primers are listed in (Supplemental Table S1E). Expression data were normalized by subtracting the mean CT value of the reference gene from the mean CT value of the gene of interest (ΔCT). The determination of statistical significance was carried out using the student’s t-test.

### Constructs

The Gateway recombination compatible C terminal GFP fusion destination modular vector pMod_A218-pCmYLCV:R1-CCDB-R2-GFP:HspT was made in the laboratory. Briefly, the CmYLCV promoter amplicon from pMod_B2103 vector, R1-CCDB-R2 amplicon from pEarleyGate 100 vector, and GFP-HSPTerminator amplicon from pMod_C3003 vector were amplified by PCR and assembled by using NEB HiFi assembly master mix (E2621S, NEB, USA). Later, by employing gateway cloning (www.lifetechnologies.com) *GmNIP2-1, GmNIP2-2, GmHiSil2c* (genes synthesized from Twist Bioscience, USA) genes were cloned into Pd221 entry vector by using BP-clonase enzyme. Next entry clones were moved to GFP fusion destination modular vector pMod_A218 by using LR-clonase enzyme. All the constructs were sequence confirmed by sequencing whole plasmids by Oxford Nanopore Technologies from Plasmidsaurus, Eugene, USA.

### Protoplast preparation and transformation

Tobacco protoplasts were isolated from the leaves of *Nicotiana benthamiana* and transiently transformed as described by (Yoo et al., 2007; Nwoko et al., 2023). We used 2*10^5^ protoplasts per transfection reaction, Equal concentration (15μg) of plasmids were used for the localization experiments.

### Laser Scanning Confocal Microscopy

Protoplasts, expressing fluorescent constructs transiently, were observed using a laser scanning confocal microscope (Olympus) within their culture medium after 12h of transformation. GFP fluorescence was detected utilizing a 488 nm argon ion laser line, with emission recorded using a 500–540 nm filter set. The chlorophyll autofluorescence was detected using the fluorophore set at 647 nm and emission recorded at 650-750 nm filter set. Confocal microscope images were processed in Fiji ImageJ software (Schindelin et al., 2012).

## Conclusion/Summary

Our SnRNA-Seq analysis provides a transcriptome atlas in soybean with distinct markers for different tissue types. Si acts as master regulator element amongst all other elements in regulation of biotic stress tolerance. Here, we conclude that Si regulates multiple molecular regulatory hubs such as kinases, RNAi silencing machinery genes, immune receptors (RLPs and NLRs), pathogen defense responsive hormone (SA) signaling, biosynthesis and perception genes, pathogenesis related genes and antimicrobial compound signaling genes differently in distinct cell types to enhance biotic stress tolerance.

## Data availability statement

All the soybean gene IDs for gene names mentioned in this study are listed in Supplemental Table S1-F.

## Supplemental Information

Supplementary Figures (S1 – S10)

Supplementary Tables (S1A-F)

## Author Contributions

VD: experimental design, formal Analysis, data prediction, methodology and writing original draft; LDA: snRNA-seq analysis and writing; AOK: gene validation using protoplast; LY: snRNA-seq library preparation; ABN, VPT, AS, JM and RMS: reviewing and editing manuscript; LHE: project administration, supervision and editing manuscript; RD: methodology, data analysis and editing manuscript; GBP: Conceptualization, formal analysis, funding acquisition, supervision and writing and editing manuscript.

## Acknowledgements

GBP is grateful to the Southern Soybean Research Program (SSRP), the United Soybean Board (2314–209-0401) and the State of Texas’ Governor’s University Research (GURI) for the research funding.

