## Supplementary Figures for "Identification of cell-type-specific response to silicon treatment in soybean leaves through single nucleus RNA-sequencing"

### Slide 1
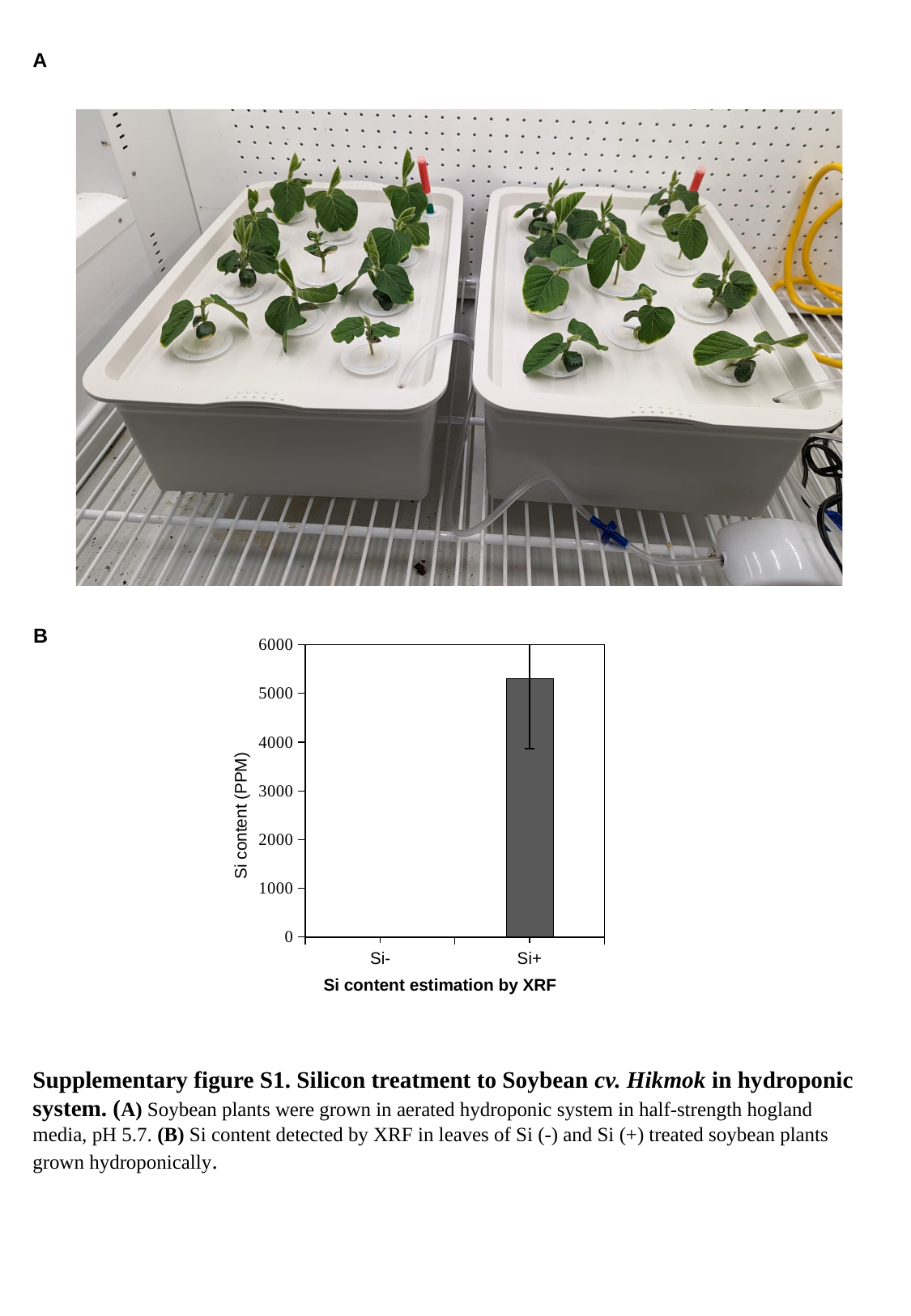

A
B
#### Chart
| Category | |
|---|---|
| Si- | 0.0 |
| Si+ | 5301.6 |Si content (PPM)
Si content estimation by XRF
Supplementary figure S1. Silicon treatment to Soybean cv. Hikmok in hydroponic system. (A) Soybean plants were grown in aerated hydroponic system in half-strength hogland media, pH 5.7. (B) Si content detected by XRF in leaves of Si (-) and Si (+) treated soybean plants grown hydroponically.

### Slide 2
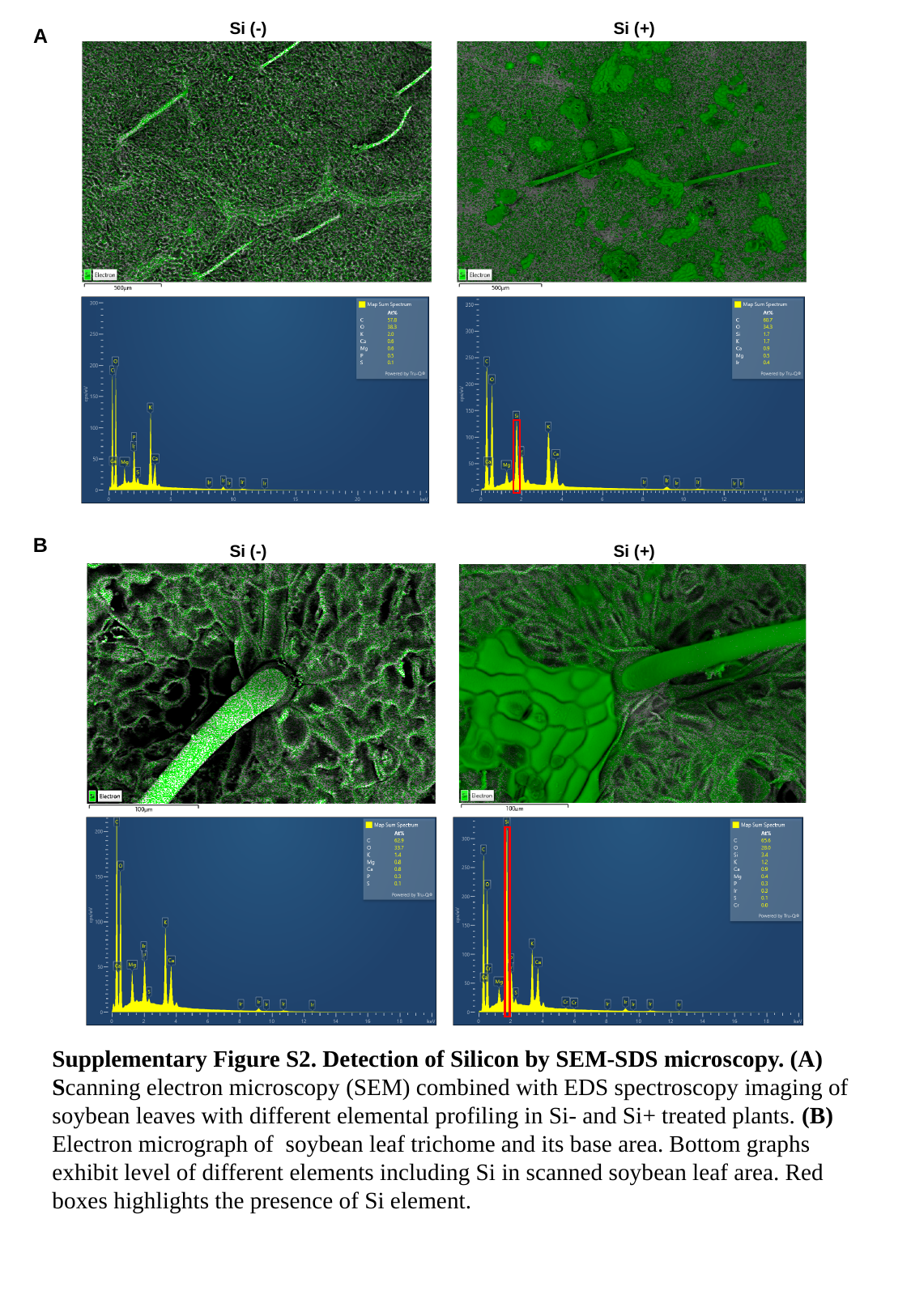

Si (-)
Si (+)
A
B
Si (-)
Si (+)
Supplementary Figure S2. Detection of Silicon by SEM-SDS microscopy. (A) Scanning electron microscopy (SEM) combined with EDS spectroscopy imaging of soybean leaves with different elemental profiling in Si- and Si+ treated plants. (B) Electron micrograph of soybean leaf trichome and its base area. Bottom graphs exhibit level of different elements including Si in scanned soybean leaf area. Red boxes highlights the presence of Si element.

### Slide 3
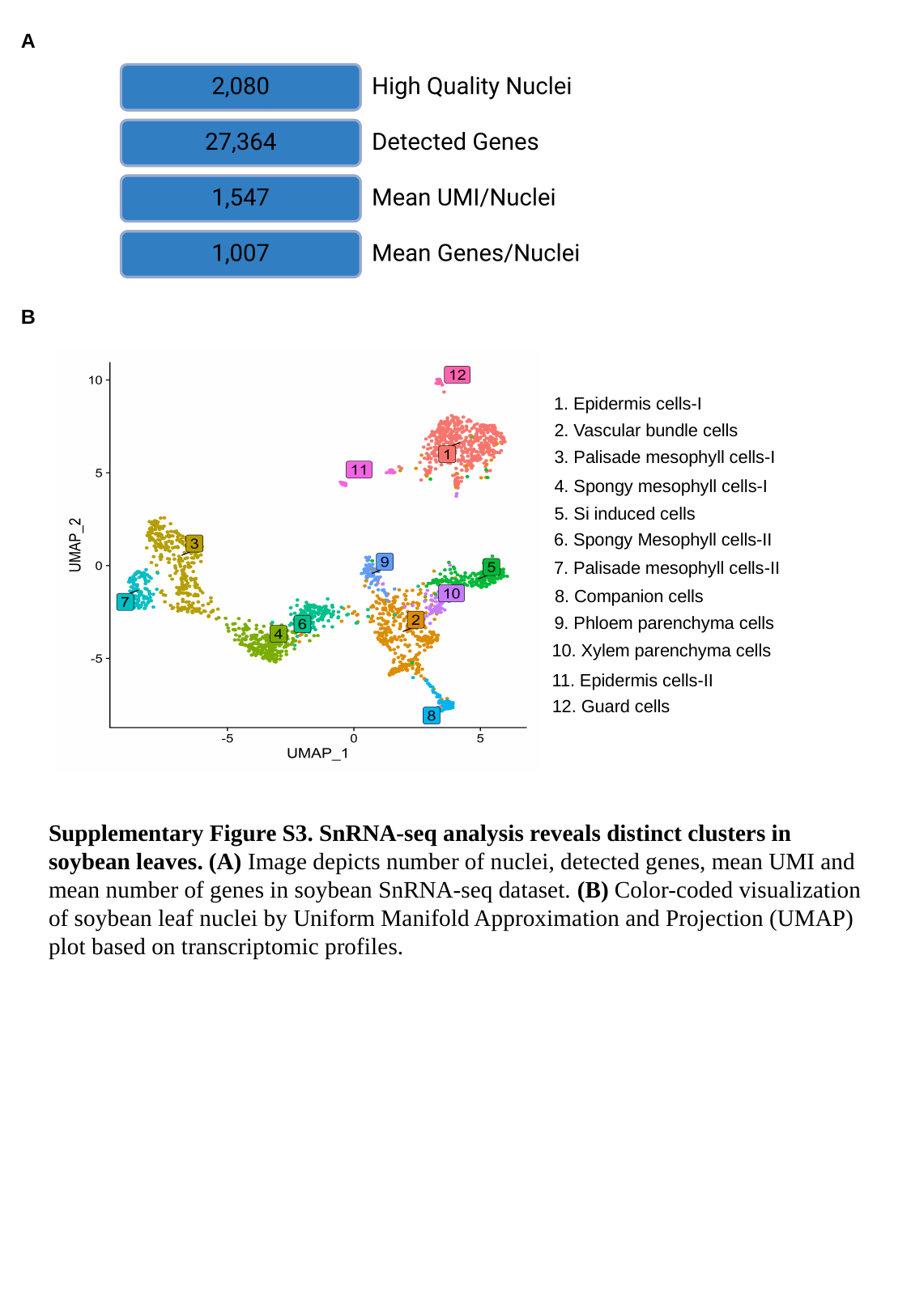

A
B
1. Epidermis cells-I
2. Vascular bundle cells
3. Palisade mesophyll cells-I
4. Spongy mesophyll cells-I
5. Si induced cells
6. Spongy Mesophyll cells-II
7. Palisade mesophyll cells-II
8. Companion cells
9. Phloem parenchyma cells
10. Xylem parenchyma cells
11. Epidermis cells-II
12. Guard cells
Supplementary Figure S3. SnRNA-seq analysis reveals distinct clusters in soybean leaves. (A) Image depicts number of nuclei, detected genes, mean UMI and mean number of genes in soybean SnRNA-seq dataset. (B) Color-coded visualization of soybean leaf nuclei by Uniform Manifold Approximation and Projection (UMAP) plot based on transcriptomic profiles.

### Slide 4
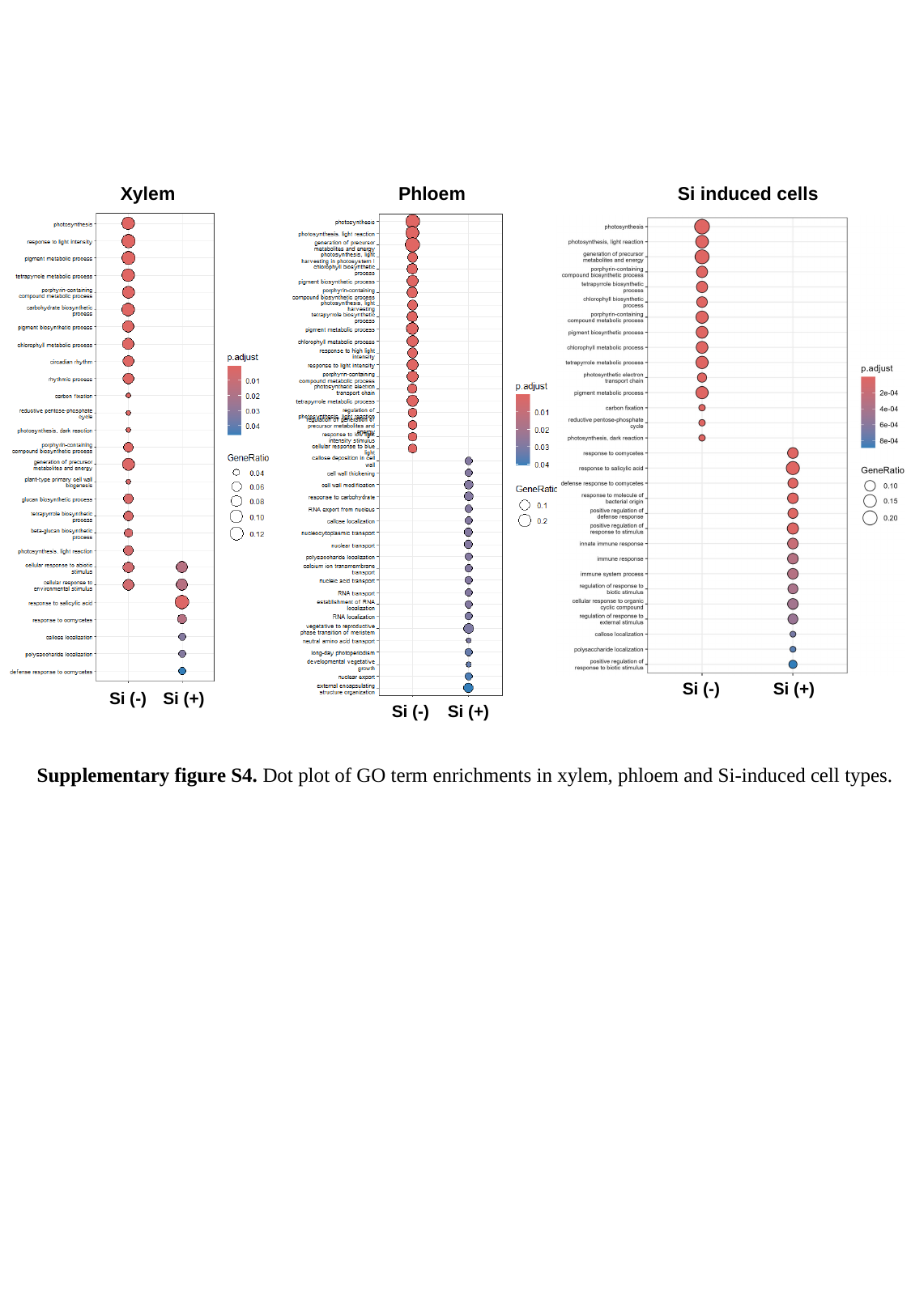

Xylem
Phloem
Si induced cells
Si (+)
Si (-)
Si (+)
Si (-)
Si (+)
Si (-)
Supplementary figure S4. Dot plot of GO term enrichments in xylem, phloem and Si-induced cell types.

### Slide 5
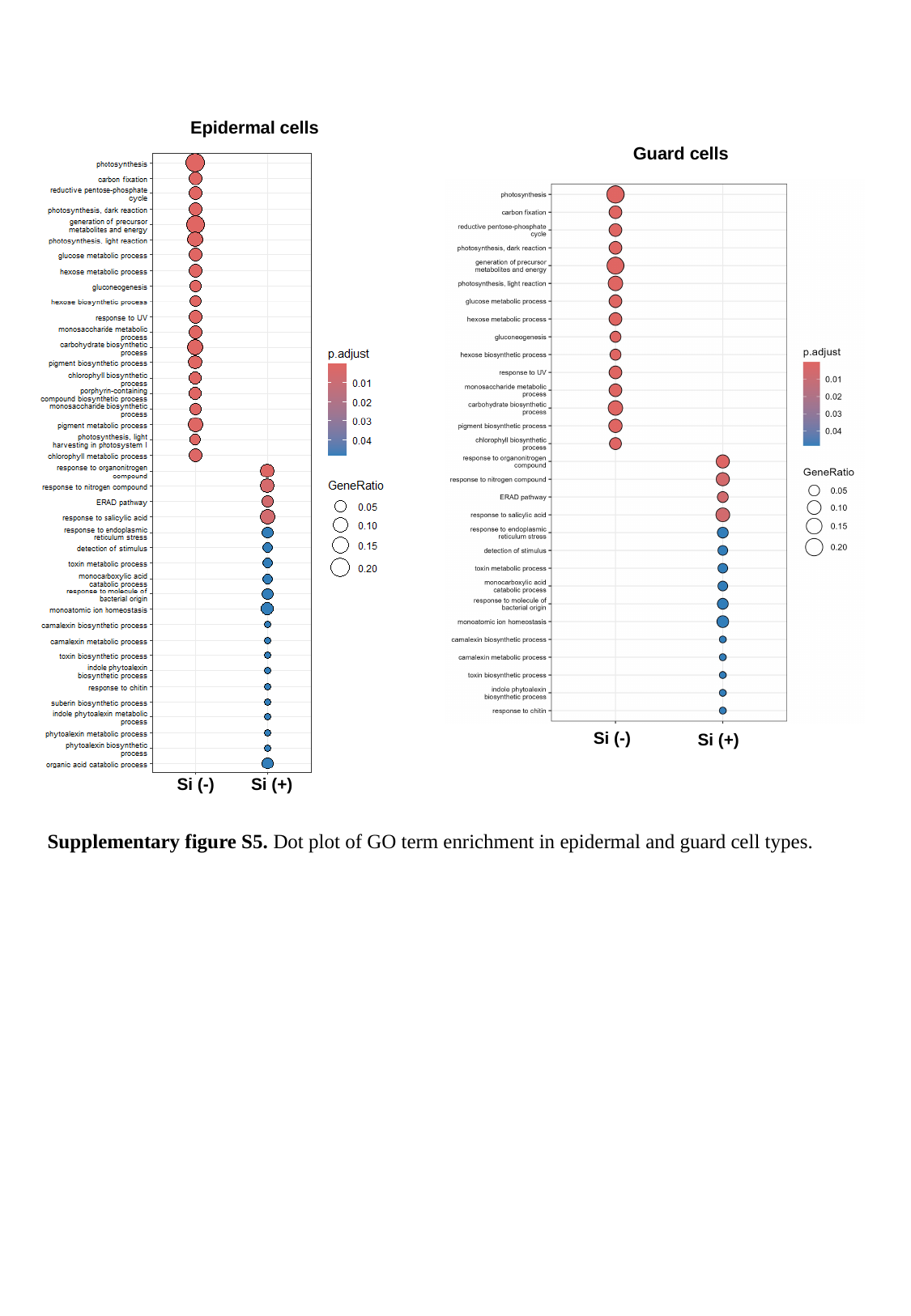

Epidermal cells
Guard cells
Si (-)
Si (+)
Si (+)
Si (-)
Supplementary figure S5. Dot plot of GO term enrichment in epidermal and guard cell types.

### Slide 6
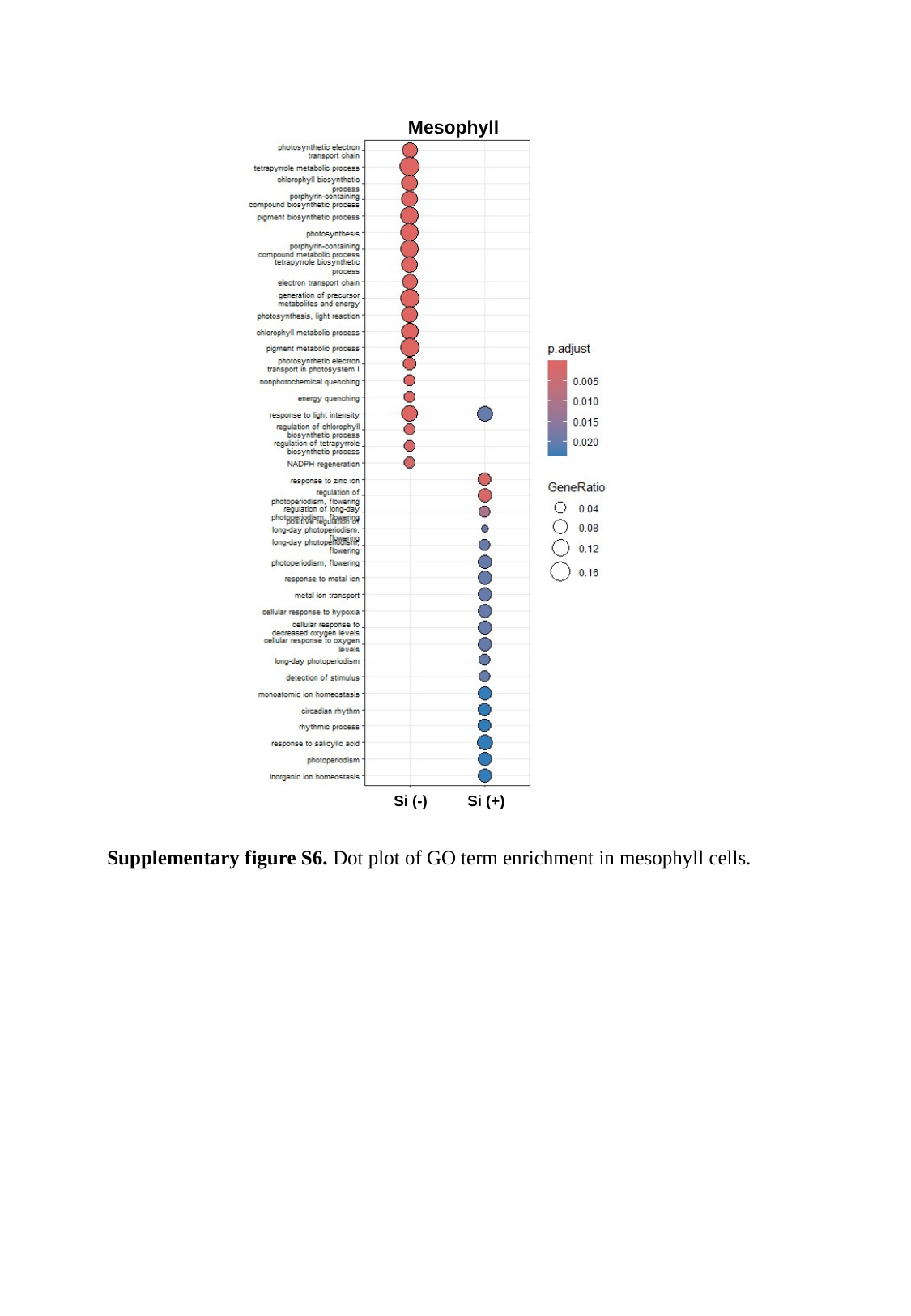

Mesophyll
Si (-)
Si (+)
Supplementary figure S6. Dot plot of GO term enrichment in mesophyll cells.

### Slide 7
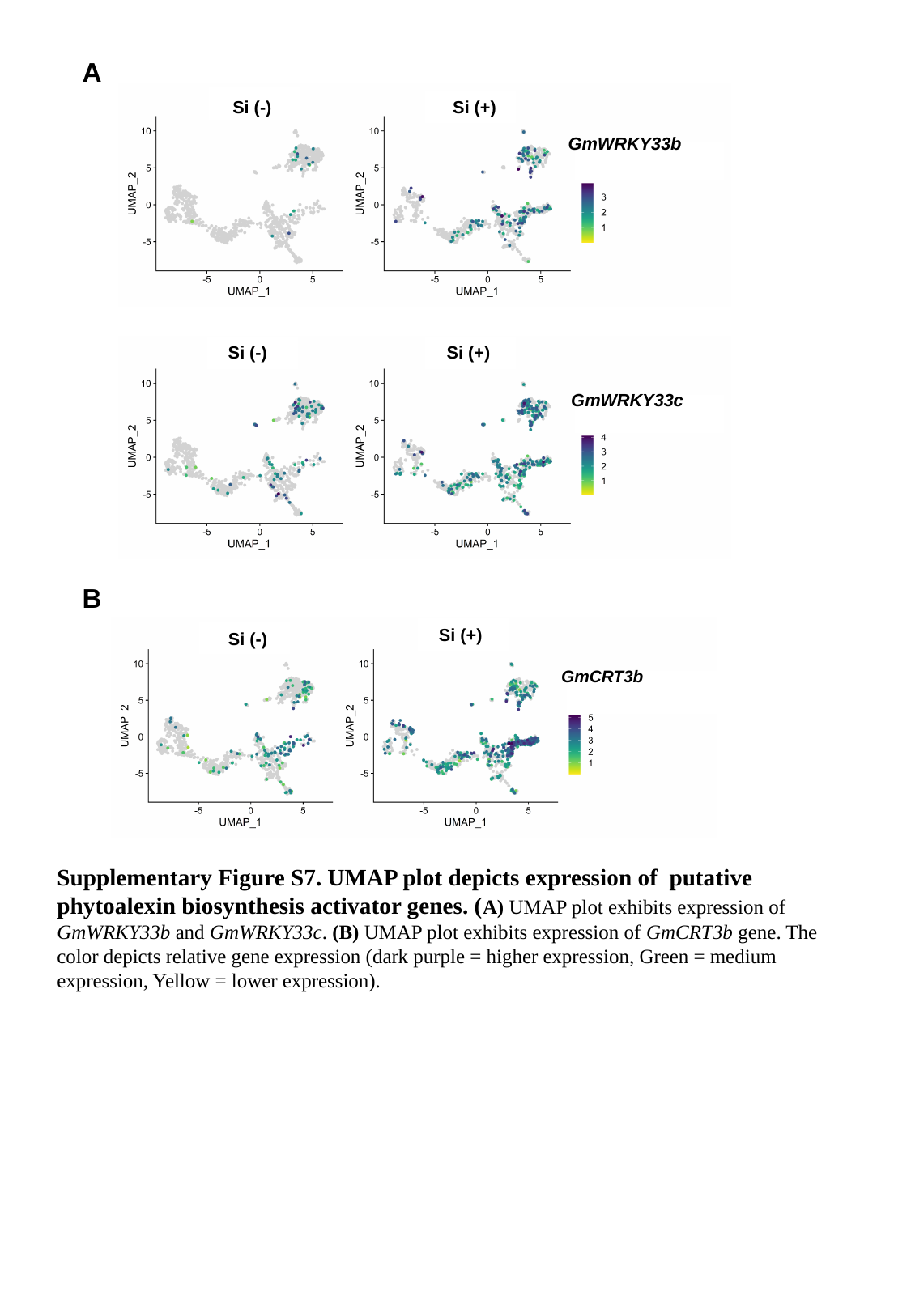

A
Si (+)
Si (-)
GmWRKY33b
Si (-)
Si (+)
GmWRKY33c
B
Si (+)
Si (-)
GmCRT3b
Supplementary Figure S7. UMAP plot depicts expression of putative phytoalexin biosynthesis activator genes. (A) UMAP plot exhibits expression of GmWRKY33b and GmWRKY33c. (B) UMAP plot exhibits expression of GmCRT3b gene. The color depicts relative gene expression (dark purple = higher expression, Green = medium expression, Yellow = lower expression).

### Slide 8
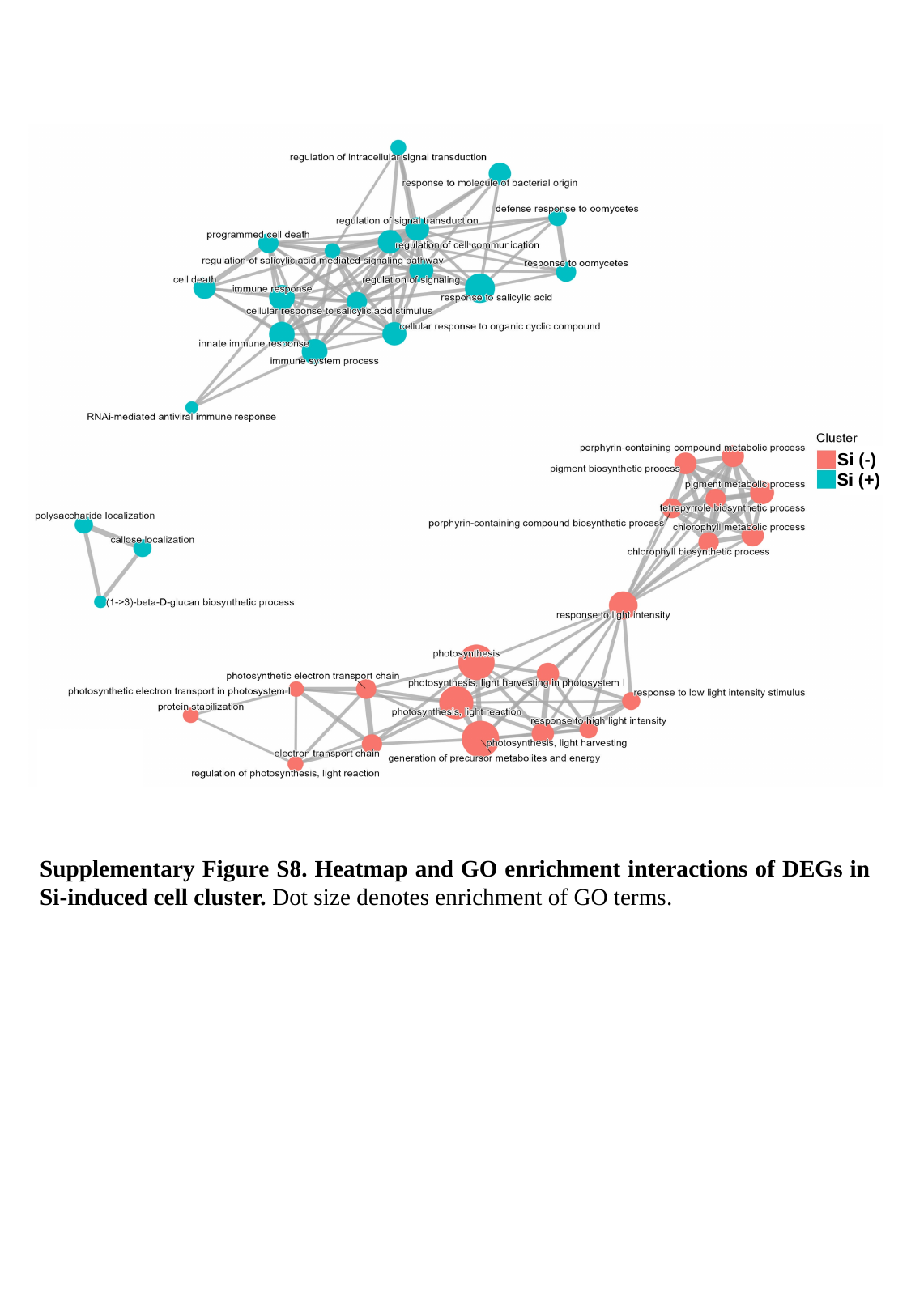

Si (-)
Si (+)
Supplementary Figure S8. Heatmap and GO enrichment interactions of DEGs in Si-induced cell cluster. Dot size denotes enrichment of GO terms.

### Slide 9
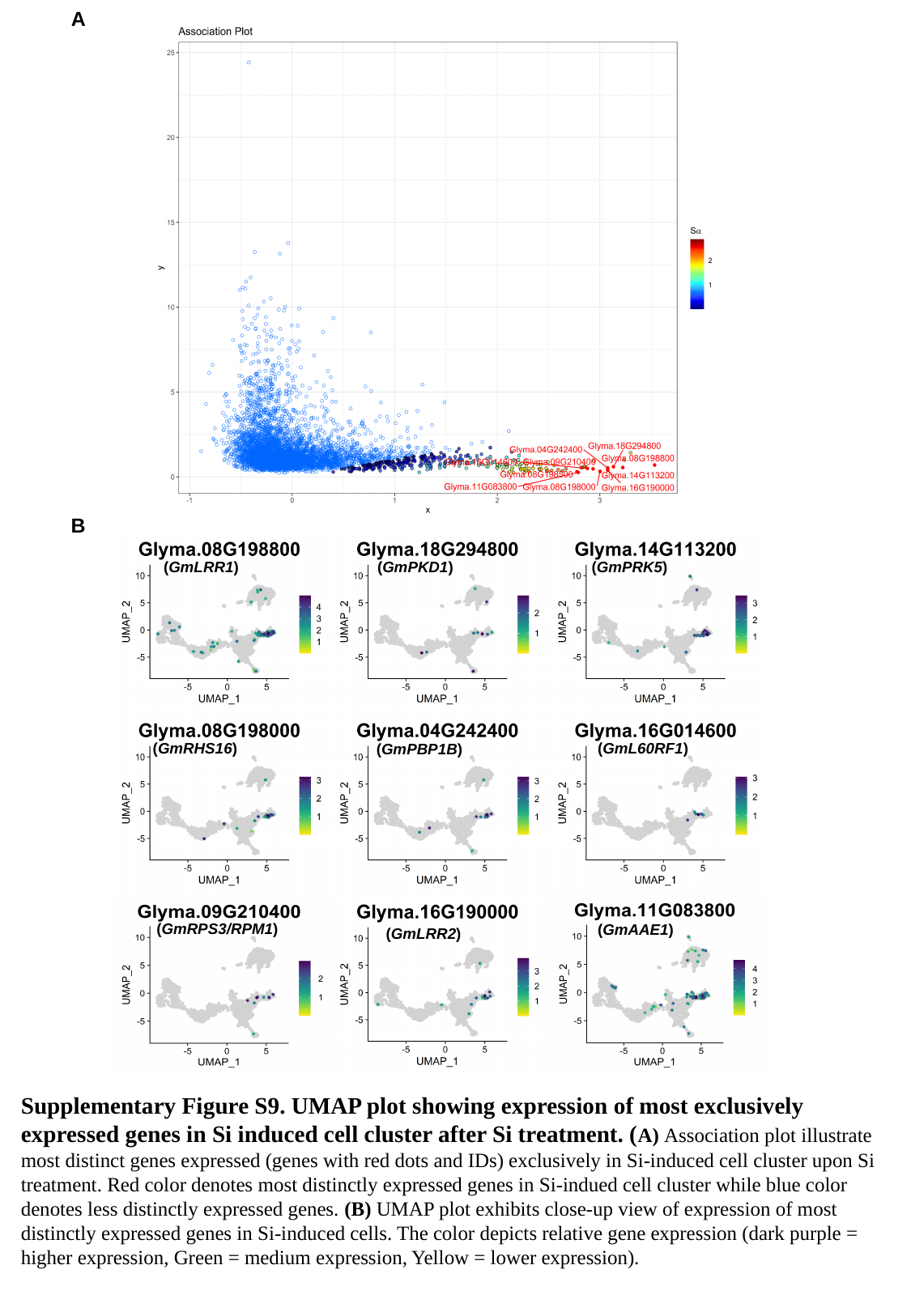

A
B
(GmLRR1)
(GmPKD1)
(GmPRK5)
(GmRHS16)
(GmL60RF1)
(GmPBP1B)
(GmRPS3/RPM1)
(GmAAE1)
(GmLRR2)
Supplementary Figure S9. UMAP plot showing expression of most exclusively expressed genes in Si induced cell cluster after Si treatment. (A) Association plot illustrate most distinct genes expressed (genes with red dots and IDs) exclusively in Si-induced cell cluster upon Si treatment. Red color denotes most distinctly expressed genes in Si-indued cell cluster while blue color denotes less distinctly expressed genes. (B) UMAP plot exhibits close-up view of expression of most distinctly expressed genes in Si-induced cells. The color depicts relative gene expression (dark purple = higher expression, Green = medium expression, Yellow = lower expression).

### Slide 10
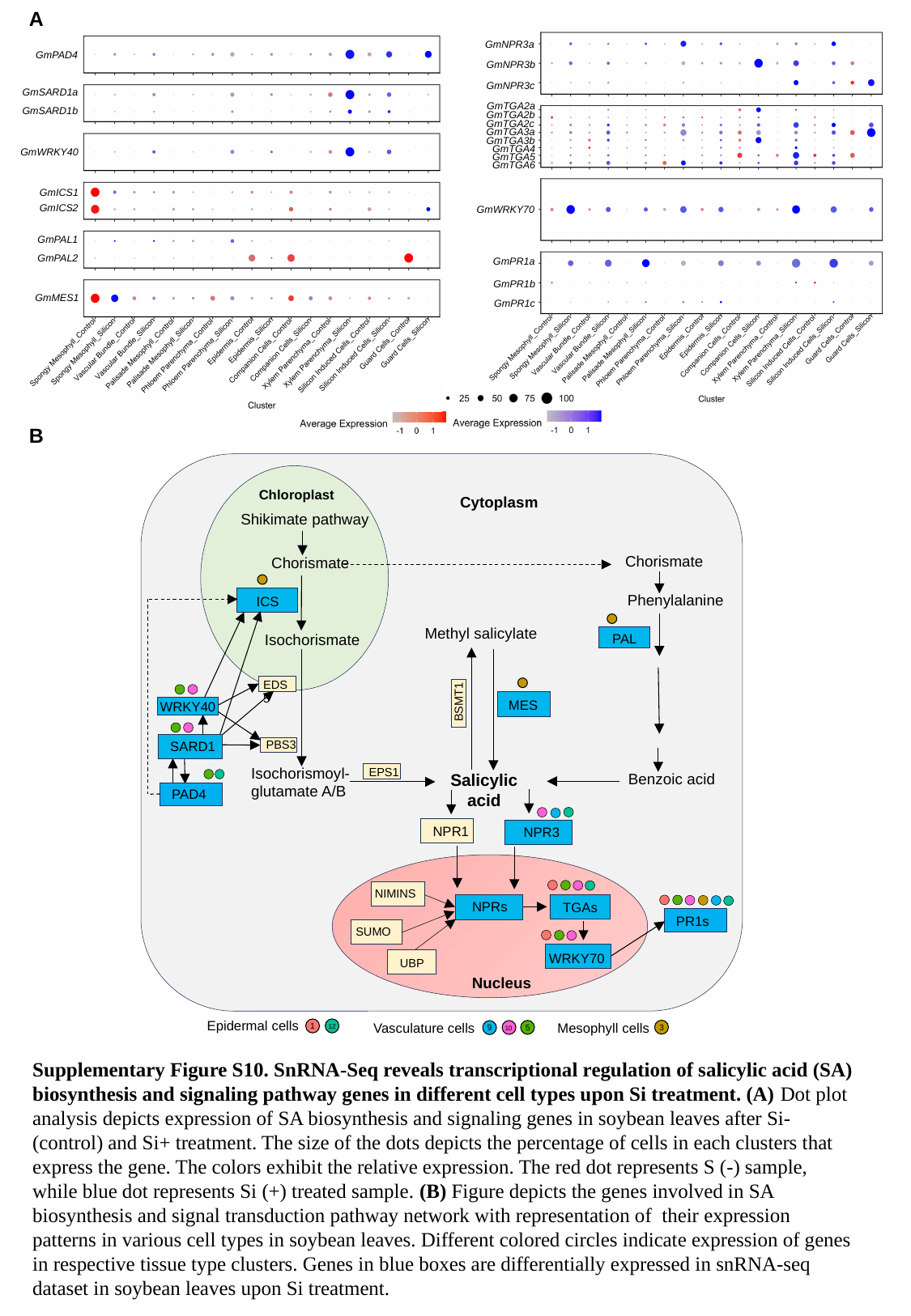

A
GmNPR3a
GmPAD4
GmNPR3b
GmNPR3c
GmSARD1a
GmTGA2a
GmSARD1b
GmTGA2b
GmTGA2c
GmTGA3a
GmTGA3b
GmTGA4
GmWRKY40
GmTGA5
GmTGA6
GmICS1
GmICS2
GmWRKY70
GmPAL1
GmPAL2
GmPR1a
GmPR1b
GmMES1
GmPR1c
B
Chloroplast
Cytoplasm
Shikimate pathway
Chorismate
Chorismate
Phenylalanine
ICS
Methyl salicylate
Isochorismate
PAL
EDS5
BSMT1
MES
WRKY40
PBS3
SARD1
Isochorismoyl-
glutamate A/B
EPS1
Benzoic acid
Salicylic acid
PAD4
NPR1
NPR3
NIMINS
NPRs
TGAs
PR1s
SUMO
WRKY70
UBP
Nucleus
Epidermal cells
12
1
Vasculature cells
10
5
9
Mesophyll cells
3
Supplementary Figure S10. SnRNA-Seq reveals transcriptional regulation of salicylic acid (SA) biosynthesis and signaling pathway genes in different cell types upon Si treatment. (A) Dot plot analysis depicts expression of SA biosynthesis and signaling genes in soybean leaves after Si- (control) and Si+ treatment. The size of the dots depicts the percentage of cells in each clusters that express the gene. The colors exhibit the relative expression. The red dot represents S (-) sample, while blue dot represents Si (+) treated sample. (B) Figure depicts the genes involved in SA biosynthesis and signal transduction pathway network with representation of their expression patterns in various cell types in soybean leaves. Different colored circles indicate expression of genes in respective tissue type clusters. Genes in blue boxes are differentially expressed in snRNA-seq dataset in soybean leaves upon Si treatment.
